# Biophysical fitness landscape design traps viral evolution

**DOI:** 10.1101/2025.03.30.646233

**Authors:** Vaibhav Mohanty, Eugene I. Shakhnovich

## Abstract

Evolutionary adaptation is often visualized as a population’s stochastic climb toward the top of a fitness landscape. While there exist approaches to design or synthetically evolve proteins into desired structures, there is a lack of methodology for designing, tuning, and quantitatively reshaping the fitness landscapes themselves on which protein evolution takes place. Here, we introduce foundational principles of fitness landscape design (FLD) to customize the structural peaks and valleys of biophysical fitness landscapes with quantitative accuracy, offering robust control of long-term evolutionary outcomes. Our FLD algorithms use stochastic optimization of a chemically derived biophysical fitness model to consistently discover optimal antibody ensembles which force a target protein to evolve according to a user-specified target fitness landscape. We then apply FLD to suppress the fitnesses of two SARS-CoV-2 genotype neutral networks and to discover proactive vaccines that preemptively restrict escape variant fitness trajectories before they arise.

**Significance Statement:** Rapidly evolving viruses mutate to escape antibodies generated by the human immune system, leading to periodic waves of infection and death. Modern vaccine design approaches that focus on currently prevalent strains can be vulnerable to emerging escape mutations. An ideal strategy for proactive vaccine design requires not only immediate effectiveness, but also control over the viral fitness landscape to ensure optimal suppression of escape variants. Here, we introduce biophysics-based computational algorithms to discover optimal antibody ensembles that quantitatively reshape the viral fitness landscape to induce long-term suppression of fitness trajectories. These protocols, called biophysical fitness landscape design, open the door to improved pandemic preparedness via proactive vaccine, antibody, and peptide design, thinking several steps ahead of pathogen evolution.

## 1. Introduction

Fitness, in an evolutionary context, is a genotype’s (or phenotype’s) reproductive success, generally defined as a growth rate of a strain or species ^1^. A *fitness landscape* ^2^ is the relationship between the set of all genotypes and their respective fitnesses, often visualized as a mountain-like structure ^3–6^ (Figure 1A). In the 90+ years since their introduction by Sewall Wright ^2^, fitness landscapes have proven to be a powerful tool for understanding evolutionary processes: for example, evolution can be viewed as a semi-random climb towards higher fitness peaks—a stochastic survival-of-the-fittest process^3,5,7^.

**Figure 1:**
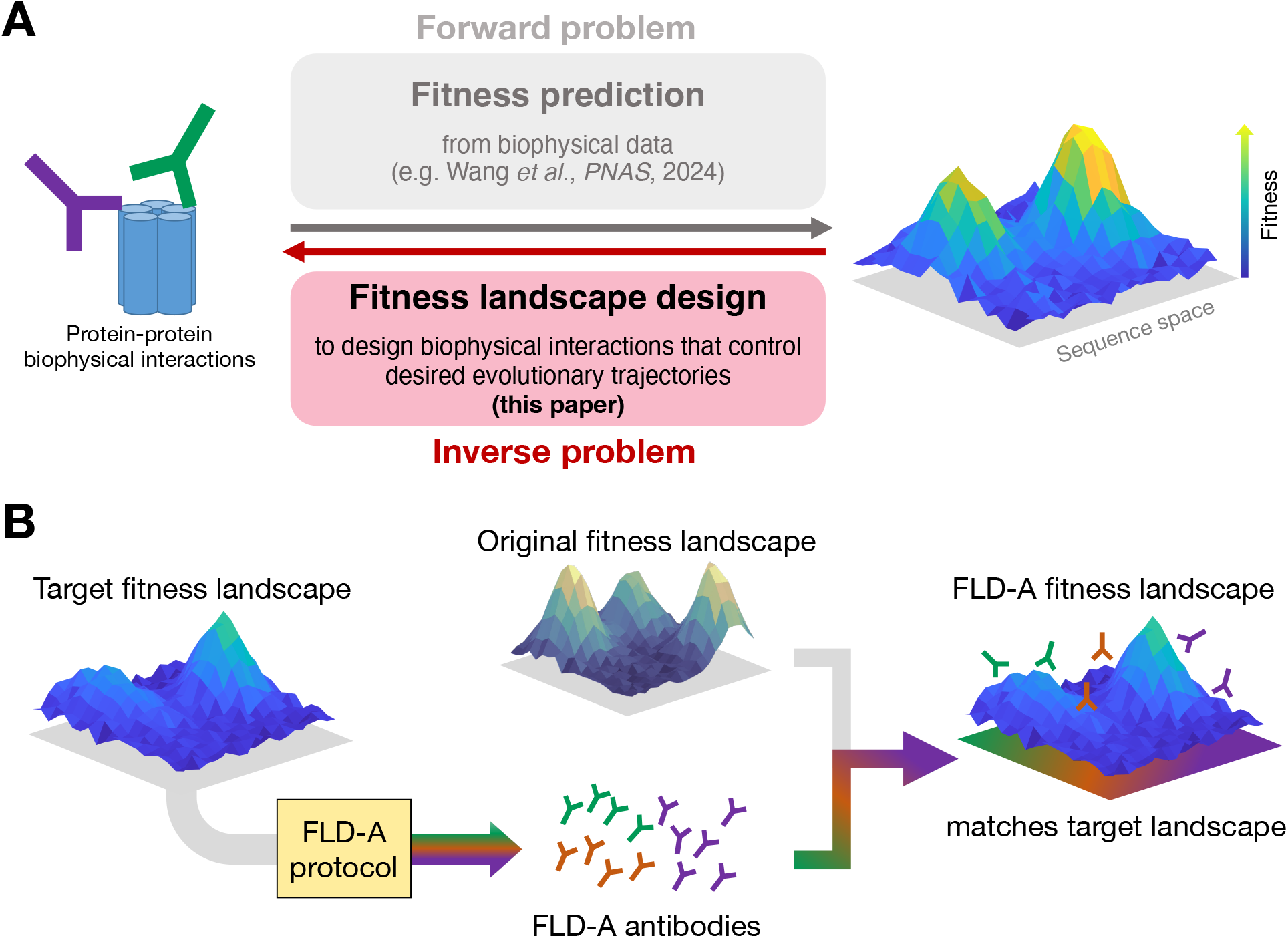
Overview of the fitness landscape design (FLD) problem, and our computational solution, FLD-A, which using antibodies to quantitatively reshape fitness landscapes. (**A**) Many studies focus on the fitness prediction *forward* problem (gray), inferring fitness landscapes from biological data, including evolutionary time series data and biophysical interaction data, including our recent work on inferring SARS-CoV-2 fitness from host-antigen and antibody-antigen binding affinities ^18^. In this paper, we introduce the *inverse* problem of *fitness landscape design* (red), where one uses a target fitness landscape to engineer biophysical interactions that influence protein evolution to take place according to the target landscape. (**B**) Our computational FLD protocols provide a solution to the fitness landscape design problem. Given a user-defined target fitness landscape for a particular target protein, the FLD-A algorithms presented in this work are used to discover an antibody ensemble which bind to the target protein and modify its experienced fitness landscape to match the target landscape. In the presence of the FLD-A antibodies, the target protein then would evolve according to the target landscape.

For proteins, it is well-established that biophysical interactions between amino acids can influence the shape of the fitness landscape and thereby affect evolutionary trajectories ^8–14^. Many studies have accordingly focused on constructing fitness landscapes by fitting theoretical fitness models to biophysical data ^6,15–18^ or directly through empirical observation of organism growth rates ^3,19,20^. Construction of fitness landscapes from data represents an important “forward” problem in evolutionary biology because it uncovers possible evolutionary paths that are enabled by nature. However, most of these studies take the fitness landscape to be externally determined and fixed, focusing instead on how evolutionary dynamics would take place on that landscape. Though there is emerging literature ^21–24^ which studies steering evolutionary paths dynamically, for example, by tuning drug dosages^23^, quantitative control over the actual shape of the fitness landscape itself is generally not well-understood.

Here, we define and introduce the corresponding “inverse” problem, which we term *fitness landscape design* (FLD): for a user-chosen target fitness landscape, how can we control protein evolution to take place according to that target landscape (Figure 1A)? For instance, how is it possible to force protein sequences *X* and *Y* to cause an organism to replicate at rates *F*_*X*_ and *F*_*Y*_, respectively? In the present work, we explore the limits of fitness landscape designability and introduce computational protocols to solve the FLD problem by focusing on viral surface proteins and the design of fitness landscapes using antibodies that can suppress long-term viral fitness gains, offering solutions to open challenges that currently prevent proactive vaccine design.

Viruses constantly face pressure from human adaptive immunity to mutate, allowing the emergence of escape variants capable of increased replication and transmission, ultimately causing a greater number of human infections and deaths^25–28^. Even if vaccination efforts are rapid—like the development of the first SARS-CoV-2 mRNA vaccine ^29^—by the time vaccines are adequately and widely deployed, viral escape mutations typically will have already emerged, seeding yet another wave of infections.

Current pandemic preparedness strategies involving vaccination are often shortsighted, focused on targeting today’s most prevalent antigens ^30–32^ but can fail to account for tomorrow’s inevitable escape mutations^33^. Approaches are emerging for the induction of broadly neutralizing antibodies (bnAbs) to target multiple strains ^34–37^ by targeting conserved peptide regions ^35–37^, for example in successful therapeutics against HIV ^38^. But, targeting of variable peptide regions in pandemic-potential viruses such as SARS-CoV-2 still remains a largely open problem. One approach might be to immunize against multiple strains, but there is an upper limit: immunizing against too many strains simultaneously can lead to B-cell “frustration” and poor immune response ^35–37^. New strategies to sequentially expose the immune system to related antigens ^37^ show promise in HIV-1^39^, but in influenza and dengue can cause “original antigenic sin” (OAS) to prevail—a phenomenon where the immune system remembers the original strain but ignores subsequent exposures to new strains, leading to outdated antibody generation and poor clinical outcomes ^40–43^.

Modern vaccine design thus continues to lag one step behind viral evolution, and waves of infection due to endemic viruses such as influenza exhibit a vicious cycle, requiring frequent and periodic updates to develop a new vaccine each year^25–28^. Developing an approach to break this cycle requires not only improved foresight into future viral evolution, but also adequate *control* over the viral fitness landscape to reshape and ultimately trap its evolutionary trajectory in a low fitness state. Proactive vaccine design against viruses thus can map onto the FLD problem, specifically the problem of designing fitness landscapes that suppress long-term fitness growth of viral escape variants.

To solve the FLD problem, we develop a set of computational methods that discover antibodies that reshape viral protein fitness landscapes for optimal suppression of escape variants, which we call *fitness landscape design with antibodies* (FLD-A). After deriving a biophysical model of viral fitness from microscopic chemical reactions, we show that a systematic choice of antibody repertoire allows different viral surface protein mutants to be assigned new fitnesses with flexibility and independence. Upon application of these antibodies, the surface proteins then evolve according to the user-defined target landscape, as validated by *in silico* serial dilution experiments using microscopic chemical reaction dynamics simulations. We then introduce algorithms to design fitness landscapes for two application tasks: (1) molding the relative fitnesses of two SARS-CoV-2 neutral genotype networks while maintaining absolute fitness reduction and (2) discovery of vaccination targets which better trap the fitness trajectory of viral escape variants compared to typical vaccination, requiring only a single target antigen and without the need for sequential vaccination.

The FLD-A protocols, summarized in Figure 1B, open the door to proactive vaccine and antibody design ^30–32,34,36,44^, offering promising new strategies for improving pandemic preparedness. Moreover, these computational methods can offer possible downstream applications to cancer therapeutics, such as reshaping cancer fitness landscapes with CAR-T cell ^45,46^ design to disfavor immune escape, and small peptide drug design ^47,48^. More broadly, the FLD-A fundamental open problem in evolutionary biology: controlling evolution to fight human disease.

## 2 Results

### 2.1 Biophysical model bridges fitness and binding affinities

#### “In vitro” fitness model for viral evolution in serial passage experiments

We first develop a biophysical model that relates a viral surface protein sequence to the growth rate that the viral strain would experience in an *in vitro* serial passage experimental setting. We derive this model analytically by starting from microscopic chemical reactions between antigen, antibodies, and host proteins and mapping them onto absolute fitness. Consider 3 sets of chemical reactions: (1) the reversible binding of a viral protein sequence **s** to a host cell receptor, (2) the reversible binding of the antigen sequence **s** to an antibody paratope sequence **a**_*n*_ which competitively inhibits antigen-host binding, and (3) the irreversible replication of the viral strain into many new copies via entry and eventual lysis of the host cell. In the Methods (and Supplementary Note 1), we show how the kinetic rate equations for these reactions combine to directly equate to the *definition* of evolutionary fitness ^1^, which then can be expressed mathematically:

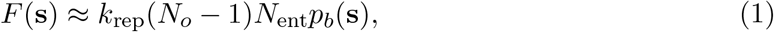

where *k*_rep_ is a single virion’s microscopic rate constant for cell entry and replication, *N*_*o*_ is the average number of offspring produced by a single virion’s replication, *N*_ent_ is the number of viral surface proteins used for host cell entry (i.e. fusion or spike proteins), and *p*_*b*_(**s**) is the probability that a single viral entry receptor with sequence **s** will be bound to the host receptors at equilibrium. This probability is given by

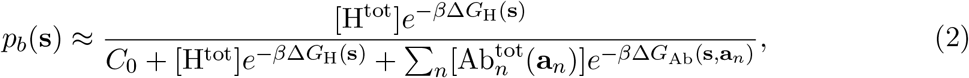

where *C*_0_ is a reference concentration to set units, *β* is inverse temperature, Δ*G*_H_(**s**) is host-antigen binding free energy, Δ*G*_Ab_(**s, a**_*n*_) is antigen-antibody binding free energy for the *n*-th antibody in the repertoire, [H^tot^] is host receptor concentration, and 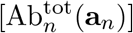 is the concentration of antibody with sequence **a**_*n*_. This model assumes that all of the relevant antibodies compete with the host receptor for binding to the viral protein’s epitope in an “*in vitro*” setting that does not include immune-mediated clearance; thus, subsequently in the paper we refer to this model as the “*in vitro* biophysical model.” The approach here is extended to include an arbitrary number of epitopes, plus immune-mediated clearance, in the following subsection.

The *in vitro* biophysical fitness function above provides a direct genotype-fitness mapping from viral antigen sequence **s** to a fitness *F*(**s**). For concreteness, for computational simulations in this paper unless otherwise stated, we focused on the biophysical fitness landscape of the SARS-CoV-2’s spike protein, for which we recently showed that a model similar to eq. (2) agrees with fitness measurements obtained from global PCR sequencing data ^18^. We considered a set of mutable loci on the receptor binding domain (RBD) of the spike. We obtained Protein Data Bank (PDB) structures of the RBD bound to the human host receptor ACE2 which facilitates viral entry ^49^ and the RBD bound to LY-CoV555, a class 2 antibody ^50^ which targets many of the same RBD amino acids involved in ACE2 binding ^51,52^. To obtain binding free energies for mutated antigens and antibodies, we allowed amino acid variation at 4 SARS-CoV-2 antigen sites and at 11 LY-CoV555 paratope sites (Figure 2A, Table S1). We performed EvoEF force field calculations ^53^ to compute host-antigen binding free energies Δ*G*_H_(**s**) which were then calibrated to experimental measurements ^54^. For antibody-antigen binding free energy prediction, a Potts model was trained ^55^ on Δ*G*_Ab_(**s, a**) force field calculations and similarly calibrated (see Methods for details). These binding free energies were plugged directly into eq. (1) to obtain fitness *F*(**s**) for SARS-CoV-2 variant sequence **s** in the presence of any chosen antibody ensemble.

**Figure 2:**
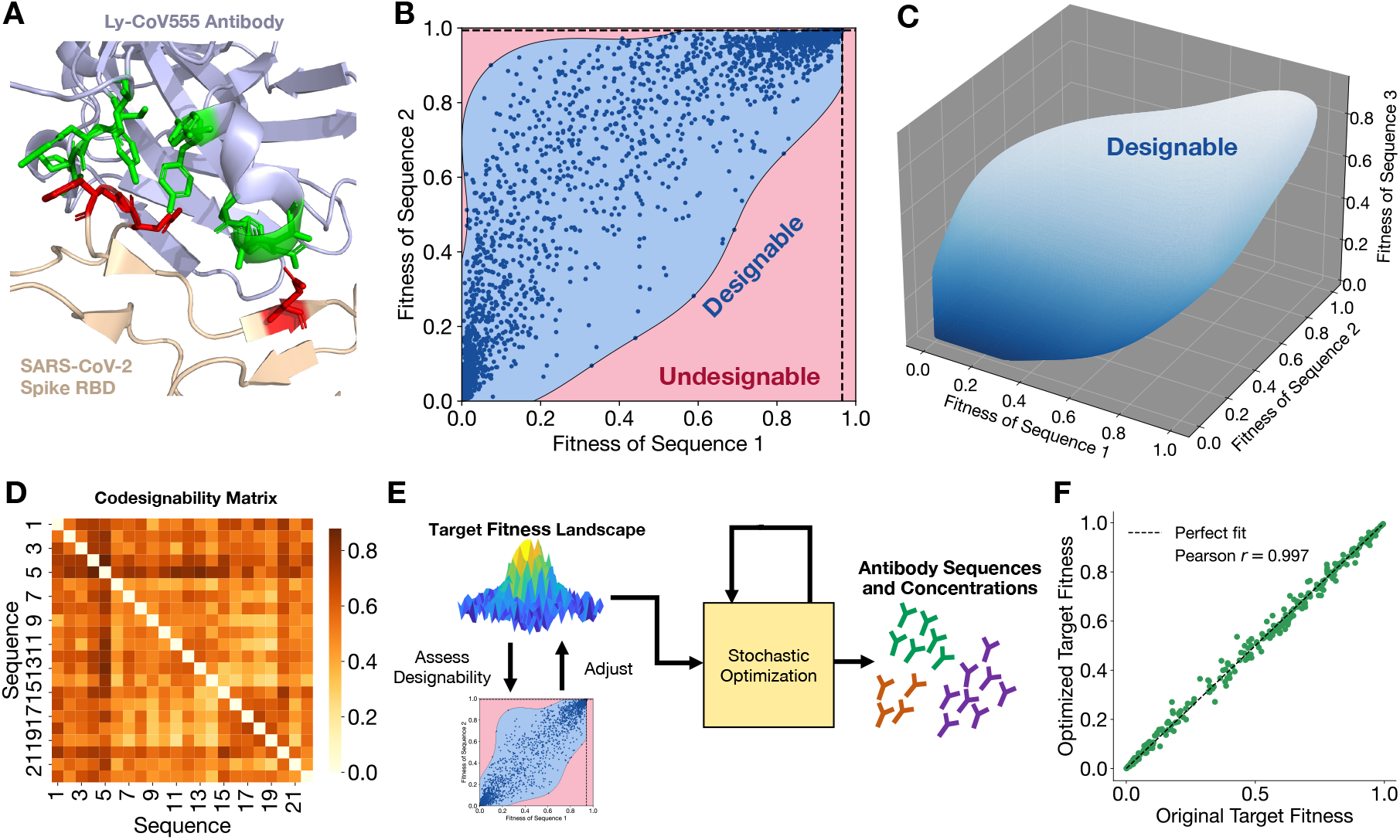
Fitness landscapes are designable and can be custom designed with stochastic optimization. (**A**) Antigen-antibody complex between SARS-CoV-2 RBD and LY-CoV555, with residues subject to variation in this study for FLD highlighted in red and green, respectively. (**B**) Designability phase diagram for two antigen sequences, obtained by sampling random antibody ensembles and using an SVM to compute the designable region. Dark blue dots are the 10,000 randomly sampled fitness landscapes, each from a randomly generated antibody ensemble. The designable (light blue) region, obtained with an SVM, indicates possible fitness values that can be physically realized by some antibody ensemble, while the red region indicates inaccessible fitness assignments. Black dashed lines indicate the maximum possible fitness for each sequence, limited by host-antigen binding affinity, according to eq. (1). (**C**) 3D designability phase diagram for three antigen sequences, also obtained by sampling 10,000 random antibody ensembles and using an SVM to compute the designable region. (**D**) Codesignability matrix for a set of sequences, with higher codesignability scores indicating a larger designable region in the 2D phase diagram exemplified by (B). (**E**) Schematic for oFLD-A protocol, which outputs the antibody sequences and concentrations which realize the target fitness landscape. (**F**) Near-exact agreement between target fitness landscape and the fitness landscape realized by antibodies discovered by the oFLD-A protocol, with each green dot representing one of 256 different epitope sequences. *Note:* (**B**), (**C**), and (**F**) use normalized fitness, scaled down by *k*_rep_(*N*_*o*_ − 1)*N*_ent_.

#### “In vivo” fitness model incorporating immune-mediated clearance and multiple epitopes

The chemistry-based approach for deriving the *in vitro* biophysical model can also be extended to include multiple epitopes, or antibody binding sites. A subset of antibodies may bind to the same site as the host receptor, thereby serving as direct competitive inhibitors of host-binding. Another subset of antibodies may compete with each other for binding to other epitopes, but they do not necessarily interfere with host-binding^56^. Macrophages and other phagocytotic cells can detect the Fc region of the virus-bound antibodies and remove the virus from free circulation in a process called Fc*γ*R-mediated clearance. By adding a fourth type of chemical reaction to the *in vitro* chemical reactions in the previous subsection to capture this immune-mediated clearance (see Methods and Supplementary Note 3), we can derive a generalized “*in vivo* biophysical fitness” function

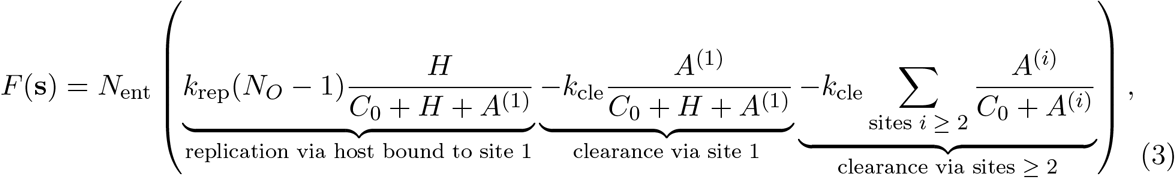

with host and epitope (site) *i*-binding antibody Boltzmann weights given by

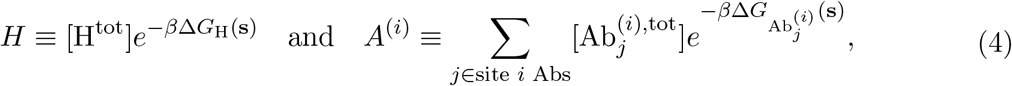

where antibody *j* which binds to epitope *i* is denoted 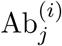. *C*_0_ is still a reference concentration, *k*_rep_ is the chemical reaction rate constant for replication, *k*_cle_ is the chemical reaction rate constant for immune-mediated clearance, and *N*_0_ is the number of offspring produced per replication event. Above, we assign “site 1,” without loss of generality, to be the epitope where the host receptor can also bind, in addition to the “site 1 antibodies.” Sites 2 and above represent distinct, non-overlapping epitopes where other antibodies can bind, but these antibodies do not impact host-binding. Thus, antibodies binding to site 1 can directly reduce cell entry probability and viral replication by competing with the host-binding (just like in the *in vitro* biophysical model), and they can additionally contribute to immune-mediated clearance, which has chemical reaction rate constant *k*_cle_. In contrast, antibodies binding to sites 2 and above do not impact cell entry and replication rate, but still contribute to immune-mediated clearance.

In order to focus on the simplest features of the biophysical model and its utility in FLD, we use the *in vitro* biophysical model for all downstream applications in this paper, unless otherwise indicated. At the end of the Results section, however, we discuss how the *in vivo* model may be paired with population-level serological data to work toward vaccine design-related applications of FLD.

### 2.2 Fitness landscapes are designable

The biophysical models provide easily computable genotype-fitness maps. Using the *in vitro* biophysical fitness model, we assessed the designability of the fitness landscape: for instance, if genotype 1 has low fitness, how low or high can genotype 2’s fitness be? For the simplest case of two randomly chosen SARS-CoV-2 antigen sequences, the space of all possible fitness assignments is shown in Figure 2B. By generating random ensembles of antibodies (see Methods), we discovered what subset of fitness assignments are practically realizable by some antibody repertoire (the blue, “designable” region in Figure 2B outlined using a support vector machine, or SVM), and what fitness assignments are not realizable by any antibody repertoire (the red, “undesignable” region). Together, the two regions produce a “phase diagram” for fitness landscape designability. For a pair of sequences, when the designable region’s area—which we call the *codesignability score*—is larger, it means that two genotypes’ fitnesses are more independent of each other, so there is a broader range of fitness choices for those sequences.

To provide a more intuitive explanation of the FLD phase diagram, consider the case when there are zero antibodies: The fitnesses of both variants will reach their maximal values (the top right of the phase diagram) because there is no neutralization. Conversely, if the viral broth culture is flooded with antibodies, both fitnesses will be as low as possible (the bottom left corner). Highly selective antibodies for one or the other strain will lead to datapoints in the top left or bottom right corners of the phase diagram, expanding the width of the designable region. Selectivity, and therefore the size of the designable region (i.e. the codesignability score), will depend on how chemically similar the antigen strains are; strains which have highly correlated binding affinities will also have highly correlated fitness assignments and therefore a narrower designable region ^57^. Thus, having a variety of antibodies with different binding affinities as well as selectivity to strains would allow for differing fitness assignments to different strains, which may be optimal in downstream tasks such as preventing certain dangerous mutations, as we will discuss in Section 2.4. The phase diagram concept can certainly be generalized to more than two sequences (Figure 2C), though visualizations become difficult beyond three dimensions. See ref. ^57^ for further discussion of FLD phase diagrams, including an analytical theory for their expected shape as well as experimental validation with an influenza dataset.

The codesignability matrix (Figure 2D) captures codesignability scores for all pairs of sequences in a set, which lets us see which sequences have more flexibility in fitness assignment. We will show in a subsequent application that the codesignability matrix can capture useful biophysical information about neutral genotype networks.

Since codesignability scores are generally greater than zero, we can manually reshape fitness landscapes for a set of antigenic sequences by finding an appropriate antibody ensemble. Concretely, our goal is to take some user-defined target fitness landscape *F*_target_(**s**), chosen from the designable region, and find an antibody ensemble that causes the viruses to actually evolve according to that target landscape. To match the biophysical fitnesses *F*(**s**) from eq. (1) to the target fitnesses *F*_target_(**s**), we perform simulated annealing ^58^ with the Metropolis-Hastings algorithm ^59^ to update antibody sequences and concentrations, progressively minimizing the least-squares loss between the target landscape *F*_target_(**s**) and biophysical landscape *F*(**s**) by iteratively updating the antibody concentrations and paratope amino acids. The output of this optimization FLD-A (oFLD-A) protocol, summarized in Figure 2E, is a list of antibody variants and their respective concentrations which would force the viruses to experience the target fitness landscape. Although we visualize the designability phase diagrams for only two or three sequences, as an illustrate example we test the oFLD-A protocol on a set of 256 antigen sequences and find that the protocol successfully and reproducibly obtains an antibody ensemble for which biophysical fitnesses nearly perfectly match the target fitnesses (Figure 2F). Thus, we have shown that custom design of fitness landscapes is indeed possible.

### 2.3 *In vitro* experiments, epidemiological data, and *in silico* simulations support the FLD pipeline and its underlying biophysical model

Next, we sought multiple independent external sources of evidence to support the FLD pipeline and the underlying biophysical model. FLD relies on having accurate fitness predictions from the biophysical model, given a viral sequence and binding free energies. If the binding energy predictor and biophysical model are both reliable, then fitnesses of real viral strains should be predictable from eq. (1) (*in vitro*) or eq. (3) (*in vivo*). Therefore, we utilized murine norovirus 1 (MNV-1) fitness data from serial dilution experiments and SARS-CoV-2 fitness measurements from global sequencing data^60^ to test whether viruses actually experience the fitnesses predicted by the biophysical model. If so, then the biophysical model is suitable for use in the FLD pipeline.

#### In vitro evidence from MNV-1 serial passage experiments

In *in vitro* serial dilution experiments, viral strains replicate in a broth containing host cells and, if desired, a neutralizing antibody (or ensemble). As illustrated in Figure 3A, after some replication time Δ*t*_*b*_, a fraction of the viral culture is transferred to a fresh broth, and samples can be sent for sequencing to determine the abundance of viral strains at each passage. The resulting viral abundance time series data can be used to infer fitness, to see if the viruses actually experienced the fitnesses predicted by the biophysical model.

**Figure 3:**
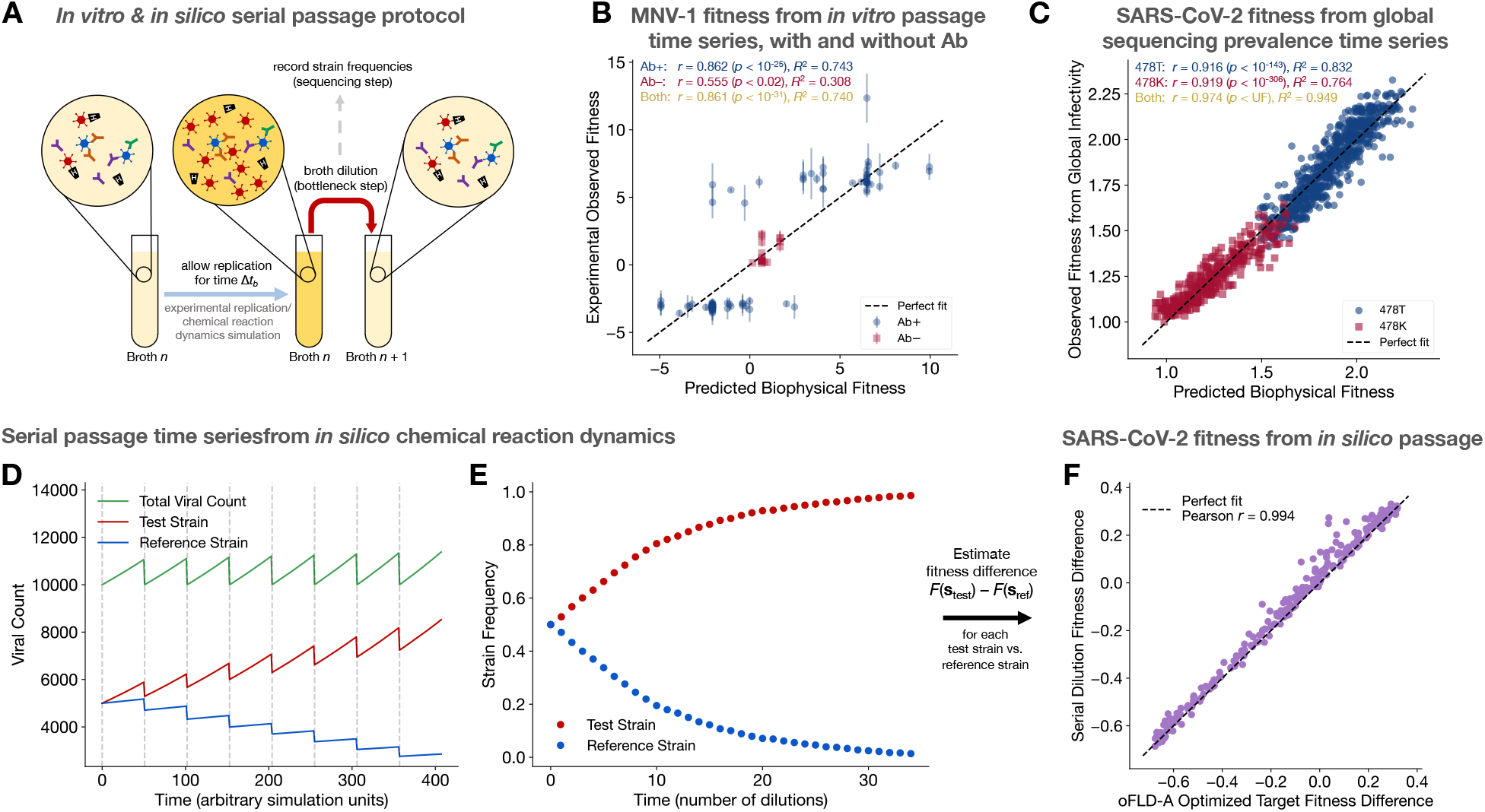
Experimental *in vitro*, epidemiological, and *in silico* viral fitness data support the FLD pipeline and its foundational biophysical model. (**A**) Schematic of *in silico* serial dilution experiments using microscopic chemical reaction dynamics. (**B**) The *in vitro* biophysical fitness model, using EvoEF force-field binding free energies, can fit empirical fitnesses estimated from time series data for evolution of MNV-1 with (Ab+) and without (Ab−) a neutralizing antibody. Additive and multiplicative prefactors are refit for each dataset while effective concentrations and free energy calibration constant are fixed across both datasets to maintain interpretability. (**C**) The *in vivo* biophysical fitness model, using experimental binding free energies from Tite-Seq, can fit empirical fitnesses estimated from global infectivity of SARS-CoV-2 variants possessing (478K) and or not possessing (478T) the T478K mutation. Additive and prefactors are refit for each dataset while the multiplicative prefactor, effective concentrations, and free energy calibration constant are fixed across both datasets to maintain interpretability.(**D**) Time evolution of viral counts from chemical reaction dynamics across multiple broth dilution steps. (**E**) Observable strain frequency data, measured at the broth dilution steps indicated by gray dashed lines in (A) and (D), representing the data available to an experimenter with no knowledge of the underlying microscopic chemical reactions. (**F**) Near-exact agreement between optimized target fitness difference (equivalent to vertical axis from Figure 2F) and simulation fitness difference. Plotted fitnesses are normalized, scaled down by *k*_rep_(*N*_*o*_ − 1)*N*_ent_.

We had previously conducted serial passage experiments using MNV-1^6^ but we had never tried to predict fitnesses of individual strains. Here, we revisited the previously published data ^6^ and extracted the time series datasets for two experimental conditions: one for evolution in the presence of monoclonal antibody A6.2^61^ (“Ab+”), and one in its absence (“Ab−”). We estimated experimental fitnesses for observed strains using the Fit-Seq2.0 software ^62^ for each condition. We then computed host-antigen and antibody-antigen binding free energies using EvoEF (the same force field used for the FLD pipeline), and substituted these energies intro the *in vitro* biophysical model, eq. (1). We then fit a single *in vitro* biophysical fitness function to both Fit-Seq2.0 datasets simultaneously, so that certain free parameters— including EvoEF calibration constants and host/antibody concentrations—could retain the same physical meaning across both experimental conditions. We allowed for the additive and multiplicative prefactors to be refit for each experimental condition, and experimental datapoints were censored if the relative error from Fit-Seq2.0 estimation exceeded a certain threshold (see Methods for fitting details).

Figure 3B shows that the *in vitro* biophysical model, using zero-shot EvoEF binding free energy predictions, simultaneously fits both experimental viral fitness datasets for a relative error cutoff 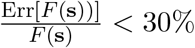 with Pearson *r* = 0.862 (*p* < 10^−25^) for the Ab+ dataset (*N* = 85 points), *r* = 0.555 (*p* < 0.02) for the Ab− dataset (*N* = 20 points), and *r* = 0.861 (*p* < 10^−31^) across both datasets combined (*N* = 85 + 20 = 105 points). See Methods and Figures S1 and S2 for additional fits for different relative error cutoffs, fits in which the multiplicative scaling prefactor was fixed across both datasets, and hyperparameter finetuning data for the simultaneous fitting of both datasets. For statistical rigor, in order to make sure the fitting power of the model was not simply due to the number of free parameters, we also independently scrambled the EvoEF binding free energy assignments Δ*G*_H_(**s**) and Δ*G*_Ab_(**s**) within each dataset and refit the models at each relative error threshold, finding that nearly all correlations were destroyed (Figure S3), thus indicating that the EvoEF force fields are indeed effective in predicting empirical fitness using the *in vitro* biophysical fitness model. Overall, these data indicate that the *in vitro* biophysical fitness model receives external support from empirical viral evolution data, even when given zero-shot force-field energies.

#### Epidemiological evidence from SARS-CoV-2 global infectivity-derived fitness

We had previously hypothesized ^18^ a phenomenological model for biophysical fitness of SARS-CoV-2 RBD mutants which is similar (but not identical) to the *in vitro* fitness model in eq. (1), but the present work is the first to show how fitness can actually be derived from microscopic chemical reaction dynamics (Supplementary Notes 1–3), providing mechanistic interpretation to the hypothesized free parameters in our previous model. Moreover, in the Methods and Supplementary Note 4 we show how, using a single-epitope approximation, the *in vivo* biophysical model—introduced here for the first time—can map onto our previously hypothesized model, up to a difference in one term which depends on the relative rates of clearance versus replication.

Now, we show that fitnesses of SARS-CoV-2 strains between wildtype and Omicron, inferred from global PCR sequencing time series data, are accurately fit by the *in vivo* biophysical model, eq. (3). Obermeyer *et al*. ^60^ had inferred empirical SARS-CoV-2 RBD mutant fitnesses using their software PyR_0_, and host-antigen plus four sets of antibody-antigen binding affinities for over 1,000 of these mutants were available from Tite-Seq ^63^ experiments done by Moulana *et al*. ^51,52^. As we did previously ^18^, the empirical fitness data were stratified into two groups: one containing mutant T478K (“478K”) and one without it (“478T”), since site 478 strongly interacts epistatically with site 614, for which binding data was not present. As with the *in vitro* MNV-1 study, we then fit a single *in vivo* biophysical fitness function to both PyR_0_ datasets simultaneously so that free parameters (Tite-Seq calibration constants and host/antibody concentrations) retain the same physical meaning across both experimental conditions. We allowed for only the additive prefactor to be refit for each experimental condition (see Methods for fitting details).

Figure 3C shows that the *in vivo* biophysical model, using Tite-Seq binding free energy predictions from Moulana *et al*. ^51,52^, simultaneously fits both experimental viral fitness datasets with Pearson *r* = 0.916 (*p* < 10^−143^) for the 478T dataset (*N* = 362 points), *r* = 0.919 (*p* < 10^−306^) for the 478K dataset (*N* = 756 points), and *r* = 0.974 (*p* value lower than machine computable limit) across both datasets combined (*N* = 362 + 756 = 1118 points). Thus, the data show that the *in vivo* biophysical fitness model receives external support from empirical viral evolution data. The improved model fitting performance relative to the MNV-1 results (Figure 3B) is likely due to the higher quality binding energy data obtained from experiment, as opposed to force field-derived energies.

#### In silico evidence from BioNetGen chemical reaction dynamics simulations

Having provided evidence for the reliability of the *in vitro* and *in vivo* biophysical fitness models, we also built an *in silico* simulation of the *in vitro* serial passage experiments in Figure 3A using the BioNetGen biochemical reaction network modeling software^64^ (see Methods for simulation details). Although the chemical reactions implemented in the software are the same microscopic binding and replication reactions previously used to build the *in vitro* biophysical fitness model, this non-equilibrium *in silico* simulation still serves as a useful ancillary test of the mathematical approximations we made in deriving the fitness model, including the separation of timescales approximation that allows for a fundamentally non-equilibrium process to be captured by a fitness that depends on equilibrium Boltzmann ratios. Furthermore, our *in silico* simulation also incorporates noise via discrete sampling of virions to populate the subsequent broth, allowing us to test how well the biophysical fitness model holds up in the face of external stochasticity not used to derive the model equations.

Returning to the 256 sequences for which we designed the fitness landscape in Section 2.2, we conducted 255 *in silico* experiments. Labeling one sequence as the reference strain, each of the remaining 255 strains separately faced the reference strain in a head-to-head competition over the course of many broth dilutions (Figure 3A). In each experiment, within a broth the two viral strains experienced exponential growth at different rates causing the total viral count to form a prototypical sawtooth-like wave ^65^ over the course of many broth dilutions (Figure 3D). At each broth dilution/transfer step, the relative frequencies of the two viral strains were recorded, akin to sequencing *in vitro* samples.

Like with the *in vitro* MNV-1 study, we extracted the fitnesses that the replicating viruses actually “experienced” by using the time series of recorded strain frequencies (Figure 3E). The fitness difference *F*(**s**_test_) −*F*(**s**_ref_) between a test strain **s**_test_ and reference strain **s**_ref_ could be extracted using a simple analytical formula ^66^ (see Methods for details). Repeating the procedure for the remaining 254 test strains provided a complete observed fitness landscape, where the observed landscape was obtained purely from genetic frequency data without knowledge of the microscopic chemistry.

In Figure 3F, we plot the observed fitness landscape versus the optimized target fitness landscape expected from the designed antibody repertoire (the vertical axis from Figure 2F). Not only do we obtain Pearson correlation *r* = 0.994, but fitness is predicted with near-exact quantitative accuracy including scaling prefactors, a merit of the biophysical fitness formula eq. (1).

Together, our *in vitro* MNV-1 data, epidemiological SARS-CoV-2 data, and *in silico* BioNetGen simulations provide compelling evidence that the biophysical models underlying FLD can be reliably used to capture the fitness landscapes “experienced” by viruses evolving in the presence of some arbitrary antibody (in the case of MNV-1) or antibodies (in the case of SARS-CoV-2), especially if given strong experimental binding free energy estimates, but even to a reasonable extent with EvoEF. We now use the *in vitro* biophysical model eq. (1) directly for applied design tasks.

### 2.4 Tuning the fitness landscape of two SARS-CoV-2 neutral genotype networks with customizable fitness penalties

We now apply the oFLD-A protocol, using the *in vitro* biophysical model, to design the fitness landscape of two SARS-CoV-2 neutral genotype networks while maintaining antiviral activity. The SARS-CoV-2 wildtype strain has 16 possible mutations at residue G485 that result in zero change in fitness. These 17 genotypes (including the wildtype) are all single mutational neighbors of each other, and are referred to as a neutral set, neutral network, or genotype network^67–70^. The Q493R mutation increases fitness of these 17 genotypes without breaking neutrality; it naturally occurred between the wildtype and Omicron variants of SARS-CoV-2^71^. Thus, the 17 Q493R− and 17 Q493R+ variants at the G485 site form two neutral networks which are interconnected fitness plateaus (Figure 4A).

**Figure 4:**
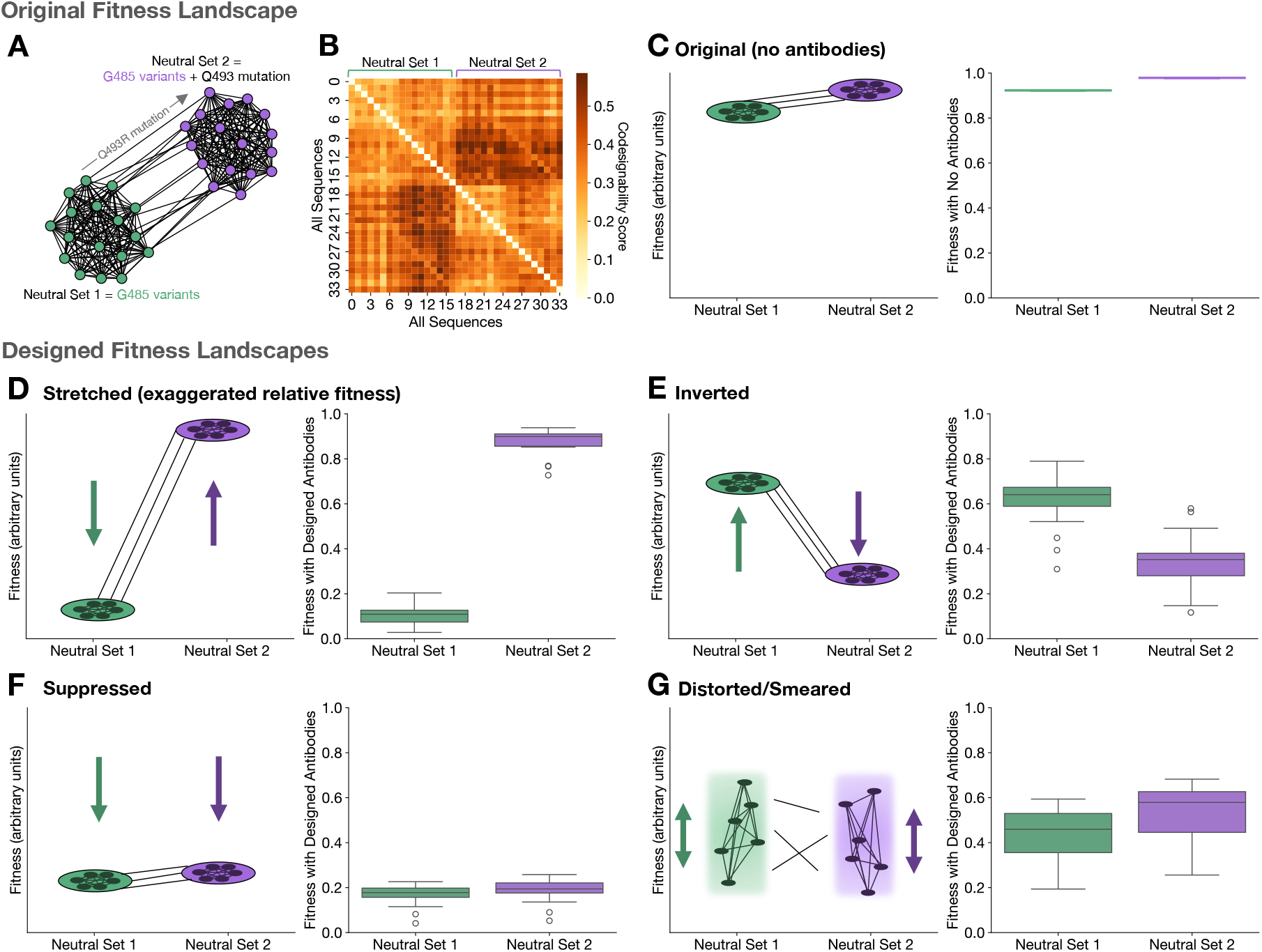
SARS-CoV-2 neutral networks are tunable. (**A**) Two SARS-CoV-2 neutral networks, each with 17 genotypes variable at residue G485, connected by the fitness-increasing mutation Q493R. (**B**) Codesignability matrix of the 34 genotypes in the 2 neutral networks, showing general trend of codesignability between neutral sets 1 and 2. (**C**) Fitnesses of original neutral networks. (**D**) Stretching relative fitnesses of the neutral sets using oFLD-A, making neutral set 2 even more fit relative to neutral set 1. (**E**) Inverting the relative fitnesses of the neutral sets using oFLD-A, turning Q493R from a fitness-increasing mutation into a deleterious one. (**F**) Suppressing the fitnesses of both neutral sets using oFLD-A, approximately maintaining their relative fitnesses but decreasing absolute fitness. (**G**) Distorting the two neutral networks using oFLD-A, breaking the neutrality of the networks altogether. Left-hand panels in (**C**-**F**) show schematics while right-hand panels show box plots of the 34 fitnesses after applying the oFLD-A protocol.

We computed the codesignability matrix for these 34 genotypes and found that the matrix separates into a 2 × 2 block matrix, with the off-diagonal blocks generally darker than the on-diagonal blocks (Figure 4B). This indicates that the two neutral networks, overall, should be codesignable as groups, and sequences within a neutral network are expected to be less codesignable. In Figure 4C, we plot a schematic representation of the two neutral networks as well as their actual fitness distributions, which are flat by definition.

Now, we demonstrate how oFLD-A can transform the original fitness landscape into a wide range of user-specified designs, while maintaining overall absolute fitness reduction. Rescaling fitnesses to a range between 0 and 1 for convenience, we set a fitness target of 0 for the Q493R− neutral set (neutral set 1) and a target of 1 for the Q493R+ neutral set (neutral set 2). The oFLD-A protocol then generates an antibody ensemble which dramatically exaggerates the relative fitness differences between the 2 neutral networks, dropping the fitness of neutral set 1 much more than the fitness of neutral set 2 (Figure 4D). By setting a fitness target of 1 for neutral set 1 and a target of 0 for neutral set 2, we can flip the relative fitnesses of the two neutral sets, causing neutral set 2 genotypes to generally have lower fitness than neutral set 1 genotypes (Figure 4E). Q493R can thus be transformed from a fitness-increasing mutation into a deleterious mutation.

By setting fitness targets to 20% of their original values, we suppress the fitnesses of both neutral sets (Figure 4F). Lastly, we can break the neutrality of both neutral sets by assigning completely random fitnesses chosen from a uniform interval to all 34 genotypes, distorting the fitness landscape entirely (Figure 4G).

FLD-A thus provides a method for molding fitness landscapes like clay: stretching, inverting, suppressing, and distorting the landscape are made possible through the optimization protocol introduced previously. We emphasize that, according to eq. (1), all of these transformations only lower—never increase—fitnesses since the maximum fitness of a strain is determined by its host receptor binding ability, which is not controlled by antibodies. This simultaneous retunability of relative fitness landscapes while suppressing absolute viral fitness makes effective and safe vaccine design possible with FLD-A.

### 2.5 Iterative FLD-A discovers proactive antibodies that trap viral fitness trajectories

We now apply FLD-A to vaccine design via antibody optimization to create evolutionary traps that restrict viral fitness trajectories within an *in vitro* context. We then discuss a path toward applying this concept on at an *in vivo*, population scale to develop improved vaccination strategies.

According to eq. (1), in the absence of any antibodies, viral fitness can increase by improving host receptor binding, mutating along the fitness landscape toward the fitness peak. After vaccinating against the wildtype or most prevalent strain, this fitness peak can shift (Figure 5A). This immune pressure encourages escape mutations, which decrease antibody-antigen binding affinity.

**Figure 5:**
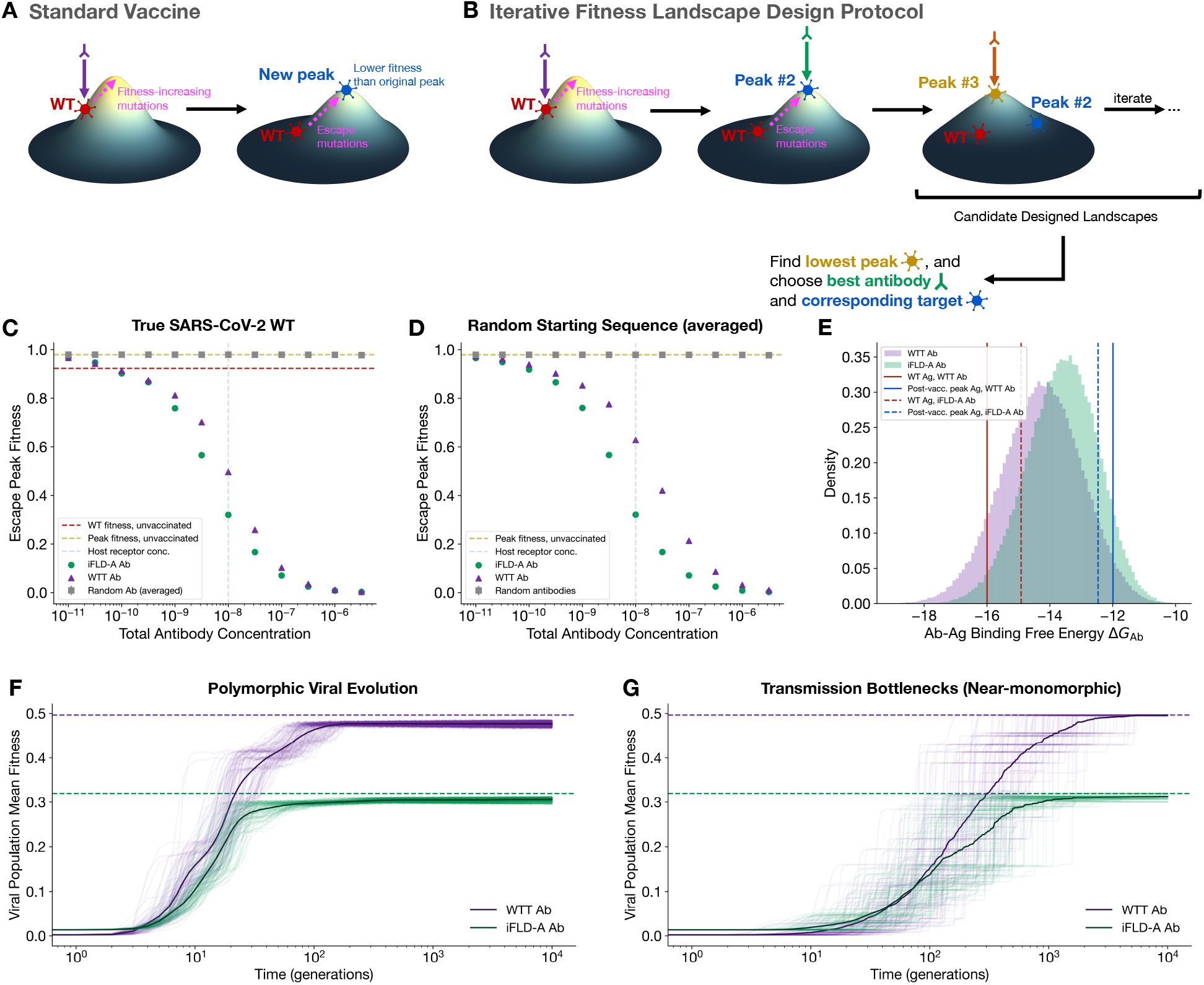
Restricting the peak fitness of viral escape variants. (**A**) Standard vaccination protocol, generating the wildtype-targeting antibody, targets the wildtype sequence. Viral escape mutations cause fitness increases towards the post-vaccination fitness peak. (**B**) iFLD-A finds the vaccine target which minimizes postvaccination peak fitness through an iterative procedure of targeting the new global fitness peak in the context of the previous antibody by searching the antigenic sequence space and computing fitness using the biophysical model in eq. (1). (**C**) Comparison of iFLD-A, standard vaccination (WTT antibody), and random antibodies, using the SARS-CoV-2 wildtype as the starting sequence. (**D**) Same as (**C**) but averaged over random starting sequences. (**E**) iFLD-A and standard vaccination antibodies’ binding affinities to both target antigens. (**F**) Polymorphic and (**G**) near-monomorphic viral evolutionary dynamics demonstrate that populations exhibit slower fitness growth and are trapped under a lower fitness ceiling by the iFLD-A antibody compared to the WTT antibody. Dashed lines indicate global maximum fitness, and translucent lines are population mean fitness trajectories from each of 100 Wright-Fisher trials.

Given the propensity for escape mutations to occur, we developed a proactive “vaccination” protocol that plans *ahead*, analogous to a chess player calculating future moves. The goal of our iterative FLD-A (iFLD-A) protocol is to find a target antigen which optimally suppresses the *post-vaccination* fitness peak (Figure 5B), so that escape mutants’ fitnesses are reduced as much as possible. Vaccine design is a complex process, since it is not clear *a priori* what specific antibodies would be generated in response to target vaccine antigen(s), at what concentrations, and towards which epitope(s). In this section, we make the simplifying approximation that, in response to “vaccination,” a single antibody is produced which is targeted toward the epitope indicated in Figure 2A which we have used in the prior oFLD-A examples throughout this paper. This “vaccine-induced antibody” is calculated by using simulated annealing to search across the antibody paratope sequence space to find a strong-binder to the target antigen, which in practice is founda at a deep local minimum or global minimum of the antibody-antigen binding free energy landscape. Though the assumption is simplifying, it can be extended to different epitopes using the *in vivo* biophysical model, and the binding affinity optimization procedure may be replaceable if sufficient population-level seroprevalence data is present, as we discuss in a later subsection, “Toward population-level proactive vaccine design.” In light of this caveat, we proceed by modeling viral evolution in the presence of an antibody as an *in vitro* process using the biophysical model from eq. (1). Although we refer to “vaccination” throughout this section, the iFLD-A protocol outlined subsequently for post-vaccination escape fitness peak suppression can be thought of as a means to design a proactive antibody which would suppress viral evolution in a setting similar to the MNV-1 experiments or BioNetGen simulations in Section 2.3.

#### Iterative FLD with antibodies (iFLD-A) protocol

We first vaccinate against the wild-type/starting strain, obtaining a high-affinity antibody (the wildtype-targeting, or WTT, antibody) to this original strain. We call this the “standard vaccination” protocol. Next, we exhaustively search over the the space of 20^4^ = 160, 000 antigenic sequences to discover the new fitness peak (“Peak #2”) in the presence of the WTT antibody, which is the escape variant with the highest fitness in the presence of the first antibody. Vaccinating against this peak escape variant generates “Antibody #2.” The peak-fitness escape variant in the presence of Antibody #2 is then targeted next. We continue to iterate this procedure, which alternates between exhaustively finding the global fitness peak and using simulated annealing to find a deep local (or global) antibody-antigen binding free energy minimum, until the new global fitness peak maps back onto a previously discovered one or until a maximum number of iterations is reached. Lastly, the protocol identifies the lowest fitness peak, which immediately reveals the ideal antibody and target antigen for fitness peak suppression (Figure 5B). Vaccinating against this target antigen would then place a low ceiling on escape variants’ fitness, generally reducing escape variant fitness more than standard vaccination would. Unlike vaccines updated *reactively* in response to newly emergent variants, restricting the fitness trajectory by vaccinating against the iFLD-A target antigen offers a *proactive* means to suppress viral proliferation.

#### iFLD-A antibody versus WTT antibody

To evaluate performance, we calculated the impact of the iFLD-A antibody, the WTT antibody, and random antibodies on the post-vaccination global fitness peak, at fixed antibody concentrations (Figure 5C). In general, the iFLD-A antibody consistently outperformed the WTT antibody and random antibodies in reducing the peak fitness of escape variants (Figure 5C; extended results in Figure S4A). To see if the same phenomenon would hold if the SARS-CoV-2 wildtype were different (i.e. if the pandemic had begun with a different starting sequence), we repeated the same study for starting sequences chosen at random. Averaging over an approximately uniform distribution of starting fitnesses (see Methods), we found similar results, with the iFLD-A antibody reducing the escape fitness peak more than the WTT antibody and random antibodies (Figure 5D). As expected, these differences in fitness reduction became less prominent in the limits of very high or very low antibody concentration, aside from small fluctations.

To understand why the iFLD-A vaccination targets generally work better than the standard vaccination target, we compared the binding free energies of the two antibodies to each protocol’s target antigens. Considering the SARS-CoV-2 wildtype case, we found that the iFLD-A antibody binds more strongly to the iFLD-A target antigen than the WTT antibody does (dashed and solid blue lines in Figure 5E; Figure S4B). The tradeoff is that the iFLD-A antibody has to sacrifice some binding affinity to the wildtype strain (dashed and solid red lines in Figure 5E; Figure S4C). Fortunately, this sacrifice is often small enough to ensure that the iFLD-A antibody still substantially suppresses wildtype fitness, often comparable to standard vaccination (Figure S4D). The iFLD-A protocol thus achieves a lower post-vaccination peak fitness with limited sacrifice of wildtype neutralization—the best of both worlds.

To ensure that the new vaccination protocol would not inadvertently lead to escape variants with tighter host-binding affinity, we calculated host binding free energies of the post-vaccination global peaks from both protocols. We observed that vaccination by either iFLD-A or WTT always forced escape variants to become weaker binders to host receptors compared to the global fitness peak with no antibodies (Figure S4E). Furthermore, since the iFLD-A antibody does not directly target the wildtype strain, we also wanted to confirm that the wildtype strain does not become an escape variant under the new vaccination protocol. Empirically, we found that for antibody concentrations ≥ 10^−9^, the iFLD-A antibody decreased peak escape fitness more than the WTT antibody ≥ 97.5% of the time, and for antibody concentrations ≤ 10^−9.5^, the iFLD-A antibody decreased peak escape fitness more than the WTT antibody between 75.5% and 93.5% of the time. Thus, with only the iFLD-A antibody, the escape peak fitness is maximally suppressed almost all of the time, and only in the worst case the iFLD-A returns the same antibody as the standard vaccination.

Finally, we simulated viral escape dynamics using the Wright-Fisher model (Methods) in the presence of the WTT antibody or the iFLD-A antibody both in the polymorphic regime (*N*_pop_*µL*_Ag_ ≫ 1), corresponding to large viral populations *N*_pop_ with high mutation rate per site *µ*^6^, and near the monomorphic regime (*N*_pop_*µL*_Ag_ ≪ 1) ^5^, representing small infective populations in transmission events. We found that in both polymorphic (Figure 5F and Figure S5A) and near-monomorphic (Figure 5G and Figure S5B) regimes, populations exhibited slower mean fitness growth and were trapped under a lower fitness ceiling by the iFLD-A antibody than the WTT antibody.

Together, these results suggest that, if the *in vitro* viral evolution model here is generalizable to viral dynamics at global scale, then at the start of the COVID-19 pandemic, vaccinating against the target strain identified by iFLD-A could have led to a lower ceiling on the fitness of escape variants, effectively trapping the fitness trajectory of escape variants below a lower fitness threshold. By vaccinating against a future global peak by anticipating escape mutations, the iFLD-A protocol generates a tighter-binding antibody to future escape variants with little sacrifice to the neutralization capacity of the wildtype or current dominant variant.

#### Toward population-level proactive vaccine design

How can the iFLD-A protcol, which uses an *in vitro* proxy for vaccine design, be extended to the population level? One approach is to define a population-averaged fitness landscape using the *in vivo* biophysical model to capture the wide variability in antibody repertoire across the human population:

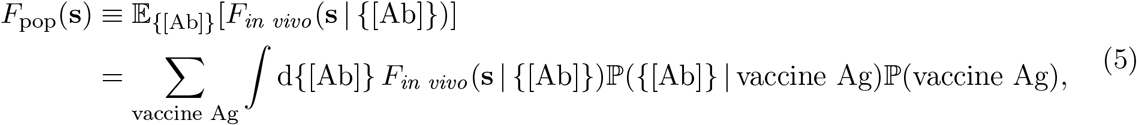

where the viral fitness landscape *F*_*in vivo*_(**s** | {[Ab]}) within an individual who has specific antibody repertoire {[Ab]} is given by the multi-epitope *in vivo* biophysical fitness function eq. (3), ℙ (vaccine Ag) is the known distribution of vaccine types across the population, and ℙ ({[Ab]} | vaccine Ag) is the the conditional distribution of antibody repertoire given which vaccine was received. We rigorously prove in Supplementary Note 6 that, assuming host receptor concentrations are roughly the same across the population, the population-average fitness is tightly bounded above and below,

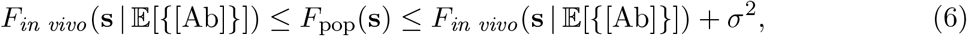

where *σ*^2^ is a weighted variance-like term for the antibody repertoire, and *F*_*in vivo*_(**s** | 𝔼 [{[Ab]}]) is simply the *in vivo* fitness for an individual who has population-averaged antibody concentrations; i.e., the population-averaged fitness is well-approximated by *in vivo* fitness with “mean-field” antibody repertoire. Retrospectively, this may explain why the microscopic *in vivo* biophysical model, with fitted concentrations, was so successful in capturing empirical SARS-CoV-2 fitnesses derived from macroscopic, population-level infectivity data in Section 2.3.

In practice, vaccine prevalence ℙ (vaccine Ag) is obtainable from surveying populations, and in this work we have introduced *F*_*in vivo*_(**s** | {[Ab]}). On the other hand, predicting the conditional distribution ℙ ({[Ab]} | vaccine Ag) is a difficult problem requiring either mechanistic modeling of within-germinal center selection ^37^ and/or machine learning models trained on large datasets. We suggest that, as calculation of *F*_pop_(**s**) becomes increasingly feasible with ongoing advancements in ℙ ({[Ab]} | vaccine Ag) prediction, it will be possible to extend the iFLD-A protocol beyond the *in vitro* setting by using eq. (5) with eq. (3) in lieu of eq. (1).

## 3 Discussion

The FLD-A algorithms represents an advance toward addressing a fundamental inverse problem in evolutionary biology: the control of evolutionary outcomes via fitness landscape design, which has wide repercussions for improving pandemic preparedness and the treatment of any rapidly evolving human disease-causing agent, including microbial pathogens and cancers. In this work, we showed how a biophysical fitness function for viral surface proteins could be used in a stochastic optimization protocol to design antibody repertoires which achieve user-defined target fitness landscapes, thus allowing quantitative control over protein evolution. The primary bottleneck for realization of FLD is being able to predict fitness of evolving viruses in the presence of antibodies, given the viral protein’s sequence. With a poor fitness predictor, FLD may not succeed, but with a high-accuracy function predictor, the fitness function can be optimized in the space of antibodies to design FLD antibodies. We thus provided *in vitro* experimental as well as epidemiological evidence to show that our biophysical fitness models, given computational or experimental binding affinity data, can achieve high predictive performance of viral fitness. In doing so, we have put forth a comprehensive pipeline to design fitness landscapes for a target protein by computationally optimizing the antibodies in which the evolution takes place for tasks such as suppressing fitness gains in viral evolution.

FLD-A offers a new perspective on vaccine design. Modern rational design of epitope-targeting vaccines relies on finding high affinity antibodies for specific prevalent antigens ^30–32^ or on generating broadly neutralizing antibodies through antigen ensembles or sequential vaccination ^34,36^. However, we show with the iFLD-A protocol that, in an *in vitro* model of viral evolution, a single antibody directed against a prudent choice of target antigen can impose post-vaccination fitness thresholds on escape variants along with wildtype neutralization. We subsequently suggeste paths forward toward population-level proactive vaccine design by relating population-level fitness landscapes to the *in vivo* fitness model derived here.

The present investigation has limitations which we aim to address with ongoing and future work. Firstly, although we have established reliability of the biophysical models underlying the FLD pipeline by showing that the *in vitro* and *in vivo* biophysical models respectively capture MNV-1 and SARS-CoV-2 fitnesses aptly in the presence of some known antibody (or ensemble), we are actively working toward building an end-to-end experimental FLD platform in which verifies the fitness landscapes imposed by newly designed antibodies. Secondly, although we have shown that the EvoEF force field energies can be successfully used in the *in vitro* biophysical function to model MNV-1 empirical fitness, performance may be significantly improved by using deep learning and large language models for binding affinity prediction^18,72,73^, possibly even without *a priori* PDB structures. Generalized binding free energy predictors are currently limited, and training high-accuracy complex-specific predictors require datasets which have free energy measurements for both antibody and antigen covariation. Currently, high-throughput datasets with either antigenic mutations^51,52^ or antibody mutations ^74^ alone exist, published datasets possessing both antibody and antigen covariation or host and antigen covariation are significantly limited. As binding affinity predictors improve, the use of EvoEF in our FLD-A protocols can be replaced with state-of-the-art counterparts. Continuous space protein sequence representations ^75^ may also be useful for gradient-optimization in lieu of simulated annealing used here. Thirdly, we the biophysical models here are mathematically tractable and “simple” because they are based on a limited set of reactions which include host-antigen binding and antibody-antigen binding. These models seem to work well for the viruses studied here including MNV-1 and SARS-CoV-2, but other nontrivial physical phenomena such as folding stability ^6^, conformation changes ^18^, and viral protein cleavage ^76^ can play important effects in viral replication and fitness. Such phenomena can be included into extended biophysical fitness models. Next, we have already discussed that the vaccine-induced immune response in the context of the iFLD-A protocol is largely simplified and that antibody repertoires are difficult to predict from knowing the vaccine target antigens. At present, it is practically complicated to implement the iFLD-A protocol at population scale without incorporation of sophisticated models^37^ of an antibody response in response to vaccine target antigens. However, we have suggested a path forward by utilizing our *in vivo* biophysical fitness model, which allows for multiple epitopes and antibody types and is mathematically related to the population-level fitness.

In addition to the future directions inspired by the limitations above, this work sets the stage for many additional avenues for applications and advancements stemming from the FLD-A protocols presented here. These include epidemiological modeling of iFLD-A vaccine effects, small peptide drug design ^47,48^, designing evolutionary fitness traps for cancers through CAR-T cell receptor design ^45,46^ or optimization of infused monoclonal antibodis, and more.

Foremost, this work introduces the concept of fitness landscape design as an inverse problem in evolutionary biology and puts forth computational methods and empirical evidence to realize FLD with antibodies. Fitness landscapes are no longer simply visual aids or theoretical tools ^3–6^ as they have been for nearly a century^2^, but rather are entities designable using biophysical principles that are capable of controlling longer-term evolutionary outcomes to fight pathogens.

## 4 Methods

### 4.1 Biophysical fitness from microscopic chemical reactions

#### In vitro biophysical fitness model

In the *in vitro* biophysical fitness model, we consider a well-mixed solution of virions, antibodies, and host cells. As in the main text, we denote the viral strain as **s**, the chemical species referring to the entire virion of that strain as V(**s**), and the chemical species referring to that strain’s surface protein (fusion or spike) responsible for host cell entry as V_ent_(**s**). Similarly, the relevant subset of the antibody sequence (the paratope) is called **a**, with the *n*-th type of antibody in the ensemble with paratope **a**_*n*_ called Ab_*n*_(**a**_*n*_). Host cell receptors are called H, concentrations of any chemical species X are denoted by the usual brackets [X], and two bound chemical species X and Y are denoted as X · Y.

The definition of absolute fitness *F*(**s**) for a viral strain **s** is the baseline replication rate of the virus

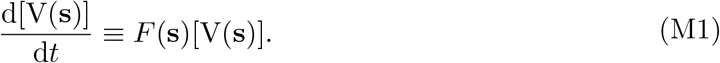

Our model starts with three chemical reactions: the host-antigen binding reaction,

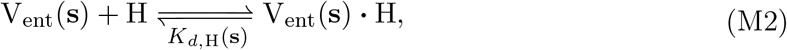

with dissociation constant *K*_*d*,H_(**s**), the antibody-antigen binding reaction

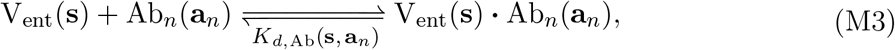

with dissociation constant *K*_*d*,Ab_(**s, a**_*n*_), and the replication reaction for a virion bound to *j* host receptors

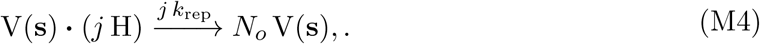

where *k*_rep_ is the baseline rate of cell entry and replication via each of the *j* receptors, and *N*_*o*_ is the number of offspring produced. From these reactions, we calculate (see Supplementary Note 1 for full derivation) that

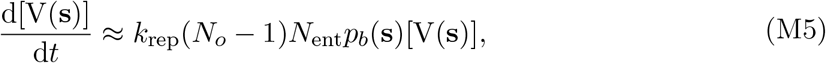

where *N*_ent_ is the number of viral entry proteins on the surface, and *p*_*b*_(**s**) is the approximate fraction of viral entry proteins which are bound to host receptors at equilibrium:

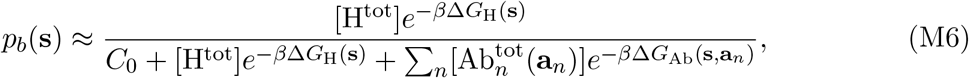

with *C*_0_ being a reference concentration to set units, *β* being inverse temperature, and Δ*G*_H_(**s**) = *β*^−1^ log *K*_*d*,H_(**s**) as well as Δ*G*_Ab_(**s, a**_*n*_) = *β*^−1^ log *K*_*d*,Ab_(**s, a**_*n*_) being binding affinities. The result from eq. (M5) is matched with the definition of absolute fitness in eq. (M1) to provide a *free energy-fitness relation*:

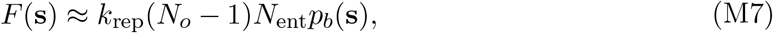

which was presented in the Results section.

#### In vivo biophysical fitness model

To extend the *in vitro* biophysical model to *in vivo* settings which include immune clearance, we must add an additional chemical reaction in addition to eq. (M2), eq. (M3), and eq. (S7):

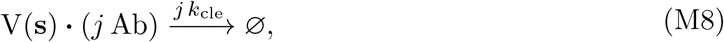

which indicates that a virion bound to *j* antibodies *of any kind* can be cleared via Fc*γ*R-mediated clearance at a rate proportional to *j* × *k*_cle_, where *k*_cle_ is the baseline clearance rate constant.

We can then carefully keep track of epitope indices as well as the individual types of chemical species (host, various antibodies) that bind to each of the epitopes and use the four types of chemical reactions to derive the *in vivo* biophysical fitness model eq. (3). A complete derivation is shown in Supplementary Note 3.

### 4.2 Fitness calculations for *in vitro* biophysical fitness model

We apply the *in vitro* biophysical fitness model, eq. (1), throughout the paper to SARS-CoV-2 to create designability phase diagrams, run the oFLD-A algorithm, run the *in silico* serial dilution experiments, and run the iFLD-A algorithms. Here, we discuss how fitness calculations are performed for these computational applications. Procedures for fitness calculations for the empirical MNV-1 and SARS-CoV-2 are discussed in Section 4.5 and Section 4.6, respectively.

#### Force field binding free energy calculations

We obtained PDB structures for the antigen-antibody complex (PDB number 7KMG) and antigen-host complex (PDB number 7DQA). 4 antigen residues, 5 antibody light chain residues, and 6 antibody heavy chain residues (Table S1) were chosen to be the mutable residues for fitness landscape design. Each selected antigen residue was involved in contact with both the antibody and the host, defined by < 8 Å distance between C_*α*_ atoms. We exhaustively computed host-antigen binding free energies Δ*G*_H_(**s**) for all 20^4^ = 160, 000 antigenic variants (i.e. all possible amino acid sequences across the 4 antigenic sites) using EvoEF force field ^53^ calculations performed at default (standard) temperature, consistent with calculation of *β* in eq. (M7) and BioNetGen simulations.

#### Potts model for antibody-antigen binding free energy

For the antigen-antibody complex, mutations could occur at the 4 antigenic sites and at all 11 antibody sites, so with 20^4+11^ ≈ 3.28 × 10^19^ possible variants, exhaustive free energy calculation is impossible. So, we randomly sampled 160,000 variants and computed Δ*G*_Ab_(**s, a**) using EvoEF. We used these data to train the parameters of a site-symmetric, spin-asymmetric chiral Potts model:

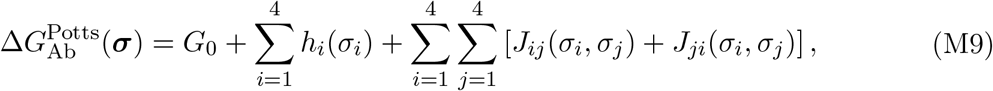

where ***σ*** is a length 15 vector combining antigen sequence **s** and antibody sequence **a**. Using a 80%-20% training-validation split, we used stochastic gradient descent to train the 90,301 parameters *G*_0_, {*h*_*i*_(*σ*_*i*_)}, and {*J*_*ij*_(*σ*_*i*_, *σ*_*j*_)}. The Potts model achieved internal consistency, with training Pearson *r* = 0.935 (Figure S7A) and validation *r* = 0.914 (Figure S7B).

#### Calibrating free energy predictions to experimental data

Starr *et al*. ^54^ conducted a deep mutational scan across the SARS-CoV-2 spike RBD and reported binding free energies. Since their mutations were not fully combinatorial, we extracted the 77 sequences in their dataset which overlapped with our antigenic sequences (19 single-site mutations at each of the 4 antigenic sites, plus the wildtype). Experimental Δ*G*_H_(**s**) measurements are plotted against the force field calculations in Figure S6A. We calibrated the force field calculations to experimental measurements by moment-matching. By computing the means *µ*_Δ*G*_ and standard deviations *σ*_Δ*G*_ of both force field and experimental overlapping datasets, we applied the linear transformation

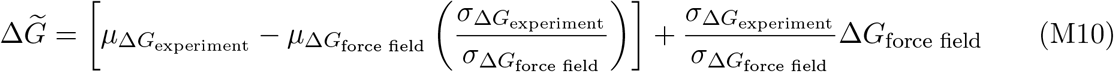

to obtain the calibrated 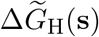 force field binding free energies (Figure S6B). This calibration allows the means and standard deviations of the overlapping data distribution to match between force field and experiment (Figure S6C). The same calibration equation was then applied to the antibody-antigen Potts model (Figure S7C and D).

#### Fitness from free energy calculations

Throughout this work, calibrated 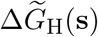 and calibrated Potts model-predicted 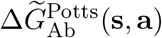 were plugged into eq. (M7) to compute fitness.

### 4.3 Computing fitness landscape designability phase diagram and codesignability matrix

To compute a fitness landscape designability phase diagram and codesignability matrix, we randomly sampled the space of antibody ensembles. Antibodies were initialized with random paratope sequences at the 11 mutable sites, and starting concentrations were initialized over a broad range of order of magnitudes, heuristically adjusting the range to ensure that all fitnesses were not all close to 0 or all close to maximal. We sampled 10,000 sets of random antibody ensembles to produce the results in Figure 2B and C, where we used 3 antibodies, and for Figure 4B, where we used 4 antibodies. For any sequence pair (or triplet), we used a support vector machine (SVM) with a radial basis function kernel to identify a decision boundary separating designable and undesignable regions. This was implemented with Scikit-learn’s^77^ OneClassSVM, with parameters *γ* = 5 (*γ* = 6 for the 3D designability diagrams) and ν = 0.0001. Codesignability scores for sequence pairs were calculated by splitting the unit square into a 100 × 100 grid on which the SVM decision was evaluated, and then summing the area of the squares where the decision was positive.

### 4.4 Stochastic optimization for fitness landscape design with antibodies (oFLD-A protocol)

The oFLD-A protocol uses a mean-square-error (MSE) loss (multiplied by a factor of 10, 000 for numerical convenience) between the biophysical fitness landscape *F*_target_(**s**) from eq. (M7) and a target fitness landscape *F*_target_(**s**):

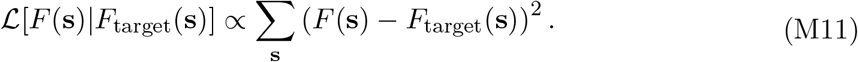

We initialized random antibody ensembles by picking random amino acids at the mutable positions. We then picked random starting concentrations which were some random number between 0 and 1 multiplied by 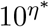, where *η*^∗^ is the exponent which minimized the loss for that starting set of random residues; for Figure 4E, we instead initialized with a remapped *η*^∗^ *↦ η*^∗^ + 1 because this tended to improve convergence to a deeper minimum.

We then perform simulated annealing, using the Metropolis-Hastings algorithm ^59^, with an exponential cooling schedule starting with simulation temperature *T*_MCMC_ = 10 and dividing by a factor of 10 after every 10,000 time steps until the 10,000 steps at the final simulation temperature (0.1 or 0.01) were completed. At each time step, we first proposed the swap of a random mutable residue of a random antibody to a random amino acid. The scaled MSE loss was recalculated, and the proposal was accepted or rejected with the standard Metropolis acceptance probability

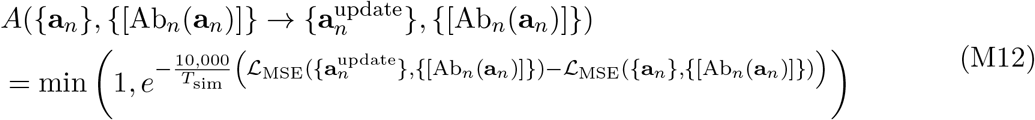

We then proposed a concentration update by randomly increasing or decreasing each concentration by 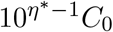 if the simulation temperature was *T*_MCMC_ ∈ {10, 1, 0.1} or by 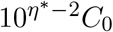 if *T*_MCMC_ = 0.01. The concentration update was accepted with probability

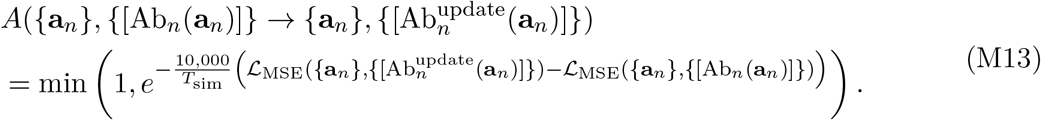

The oFLD-A algorithm thus uses alternating antibody sequence and concentration updates to find an ideal ensemble for matching the target fitness landscape.

### 4.5 Experimental MNV-1 fitness calculations and *in vitro* biophysical fitness model fitting

#### MNV-1 serial passage experiment data preparation

Time series data for MNV-1 serial passage experiments were obtained from experiment numbers 12 (bulk serial passage with no antibody, 5 timepoints) and 13 (bulk serial passage with antibody, 5 timepoints) from Rotem *et al*. ^6^. The time series data contained the read counts for each observed viral mutant, as well as number of genome copies per mL of solution. We filtered both datasets by retaining only viral mutants which had at least one read in at least two adjacent timepoints, to ensure data sufficiency for the fitness estimation software. This left *N* = 255 variants in the Ab+ dataset and *N* = 238 variants in the Ab+ dataset.

#### MNV-1 empirical fitness estimation

We then used the filtered timeseries as inputs to Fit-Seq2.0 ^62^ to obtain fitness estimates and errors for the fitness estimates. Fit-Seq2.0 asks for parameters including time between serial passages in generations. We estimated this by first noting that, at each passage, 250 *µ*L of culture were transferred to the subsequent fresh broth already containing 2 mL of media. Thus, the total culture volume was always around 2.25 mL. Under the assumption that the peak viral concentration after *g* generations in the broth was roughly the same (though it was experimentally reported ^6^ to fluctuate between 10^6^ and 10^8^), then the viral population grew to approximately 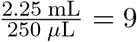 times its initial population within each culture. The number of doublings required for this growth is *g* = log_2_ 9 ≈ 3.17 generations (note that while viral burst sizes are much larger than 1 per replication event, Fit-Seq2.0 uses number of doublings as units for generations). Fit-Seq2.0 also asks for total viral population at each timepoint, which was estimated by multiplying the experimentally reported genome copies/mL by 2.25 mL. Default values were used for other parameters including half variance introduced by cell growth and cell transfer (default 1, and altering this value to 20 did not qualitatively affect fitness estimates much), optimization algorithm, and maximum number of iterations. We adjusted one line of the Fit-Seq2.0 open source code to expand the range of possible estimated selection coefficient values from *g* × [−1, 1] to *g* × [−10, 10] to allow for the possibility of strongly selected variants. Estimated fitness values were filtered in our study by calculating the relative error, which we define as the ratio 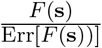 of the Fit-Seq2.0-reported fitness error to the absolute value of the estimated fitness. We filtered estimates at relative error cutoffs of 30% (Ab+ *N* = 85, Ab− *N* = 20), 50% (Ab+ *N* = 155, Ab− *N* = 25), and 100% (Ab+ *N* = 198, Ab− *N* = 49).

#### In vitro biophysical model fitting

Since Fit-Seq2.0 infers relative Malthusian fitnesses, the estimated fitnesses have some unknown additive shift. Thus, we can write the *in vitro* biophysical fitness model eq. (1) as

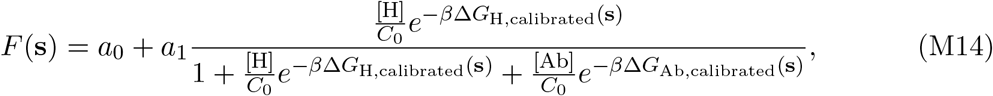

where constant *a*_0_ is additive free parameter representing the arbitrary Malthusian fitness shift and *a*_1_ = *k*_rep_(*N*_*o*_ − 1)*N*_ent_ is a multiplicative free parameter. Next, we note that EvoEF free energies—calculated from mutating PDB structures (host-antigen: 6C6Q, antibody-antigen: 7L5J)—need to be calibrated, which we implement as a linear transformation

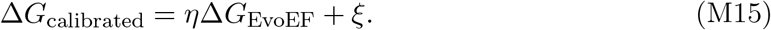

Now, defining constants *b*_0_ = *e*^−*βξ*^[H]*/C*_0_, *b*_1_ = *e*^−*βξ*^[Ab]*/C*_0_, and *c*_0_ = *βη*, we have

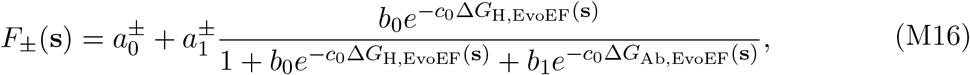

where *±* indicates different parameters for the Ab+ and Ab− datasets. To simultaneously fit both the Ab+ and Ab− datasets, allowed 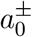 to always be fit separately for each dataset since it represents a Malthusian relative fitness shift. We also examined both the effect of allowing the multiplicative free scaling parameter 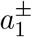 to be refit for each dataset as well as the effect of imposing a single multiplicative scaling parameter 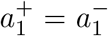 for both datasets. As expected, allowing *a*_1_ to be refit improved fitting flexibility, but both fits for both cases are reported in Figure S1 and Figure S2. The other parameters *b*_0_, *b*_1_, and *c*_0_ were fixed across the two datasets.

Simultaneous fitting was performed using `scipy.optimize.least_squares` by minimizing a hybrid least-squares loss function

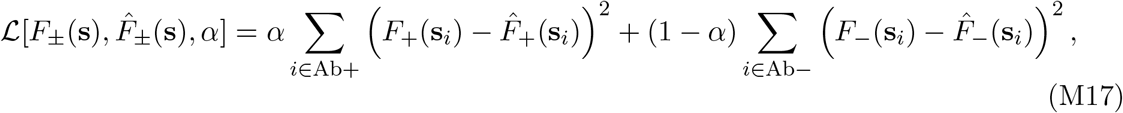

which has a hyperparameter *α* ∈ [0, 1]. For each relative error cutoff, we performed hyperparameter sweeps across this range and calculated Pearson correlation *r* and the coefficient of determination *R*^2^ for model fits and plotted the results in Figure S1 and Figure S2.

### 4.6 Empirical SARS-CoV-2 fitness calculations and *in vivo* biophysical fitness model fitting

#### SARS-CoV-2 empirical fitness and binding affinity data preparation

In a previous study ^18^, we had already extracted and filtered empirical SARS-CoV-2 fitness data from Obermeyer *et al*.’s study ^60^ for RBD mutants occurring between wildtype and Omicron BA.1. We had also extracted the corresponding host-antigen (ACE2-RBD) binding affinities from Moulana *et al*.’s study ^51^ and the antibody-antigen binding affinities for four antibodies (LY-CoV555, LY-CoV016, REGN10987, and S309) from a different study by Moulana *et al*. ^52^. Mutants were stratified into two groups (478T: *N* = 362, 478K: *N* = 756) based on the presence or absence of the T478K mutation, due to its fitness-impacting interaction with site 614 for which we did not have binding data.

#### In vivo biophysical model fitting

We show in Supplementary Note 4 that the *in vivo* biophysical model, eq. (3), can be written as

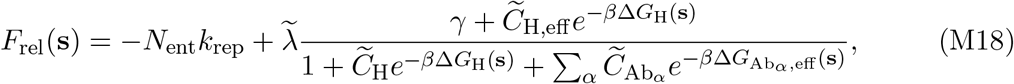

where we have defined constants 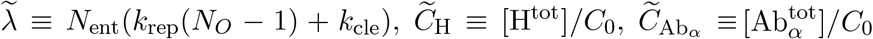, and 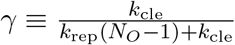.

Here, since Obermeyer *et al*.’s method PyR_0_ ^60^ inferred relative Malthusian fitnesses as a sum of additive fitness contributions from each mutation, stratifying our data by the presence of absence of the T478K means that the fitness of the two groups will simply be shifted relative to one another by an additive factor *F*_478K_ = *F*_478T_ + Δ*F*_T489K_. Again, because these are Malthusian relative fitnesses, the mutational effect Δ*F*_T489K_ can simply be absorbed into an additive free parameter capturing both some unknown Malthusian additive shifts as well as the mutational effect. Thus, we can write eq. (M18) as

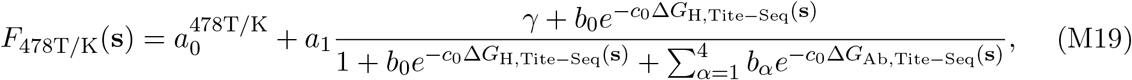

where constant *a*_0_ is additive free parameter representing a sum of the arbitrary Malthusian fitness shift from fitness estimation, as well as the additive effects of −*N*_ent_*k*_rep_ and Δ*F*_T489K_. We allow 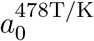 to be refit for both the 478K and 478T datasets. We have defined new constants 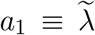 as the free multiplicative scaling parameter, 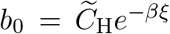 as a rescaled effective host concentration arising from linearly recalibrating Tite-Seq-derived binding free energies as done in eq. (M15) for EvoEF, 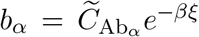 as a rescaled effective antibody concentration for each of the four antibodies *α* ∈ {1, 2, 3, 4}, and *c*_0_ = *e*^−*βη*^ as a rescaled effective inverse temperature arising from recalibrating Tite-Seq-derived binding free energies. Simultaneous fitting was performed by minimizing a hybrid least-squares loss function

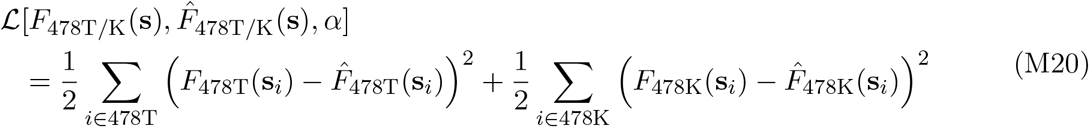

using `scipy.optimize.least_squares`. We calculated Pearson correlation *r* and the coefficient of determination *R*^2^ for model fits and plotted the results in the main text.

### 4.7 *In silico* serial dilution experiments using chemical reaction dynamics

We used the “bionetgen” Python package to generate BioNetGen ^64^ simulation files for each serial dilution experiment. For BioNetGen’s ODE simulator, concentrations use arbitrary units, as do replication rate constants; the time scale of simulations is set by the rate constants. The host concentration (or count, as the distinction is arbitrary here) was initialized at 10^9^, antibody concentrations were proportional to the ones found using the oFLD-A protocol and scaled appropriately depending on the host concentration, *N*_*o*_ was chosen to be 5, *N*_ent_ was set to 2 for computational tractability, *k*_rep_ was chosen to be 10^−11^, and the simulation time within each broth was Δ*t*_*b*_ = 3 × 10^9^. Binding dynamics needed to be faster than replication dynamics; since only dissociation constants *K*_*d*_ = *k*_off_ */k*_on_ were prescribed by the force field calculations, we chose *k*_on_ = 10^6^ with *k*_off_ = *K*_*d*_ × *k*_on_. Our chosen *k*_rep_ was appropriately much smaller than the binding rate constants.

For the first broth, viral concentrations were each initialized at 5,000. After running the BioNetGen ODE simulator for Δ*t*_*b*_, we computed viral concentrations and rounded them to the nearest integer (to approximately discretize non-integer concentrations to a viral count). We then sampled 10,000 virions, without replacement, to initialize the next broth. These viral counts, divided by the total count of 10,000, were recorded in the strain frequency timeseries. A maximum number of 50 broths were simulated for each test sequence, with broth transfers ending early if the either of the viral counts dropped below 100.

Given test strain frequency *p*(*t*) as a function of broth dilution index *t* and a total number of dilutions *t*_max_, the fitness difference *F*(**s**_test_) − *F*(**s**_reference_) was estimated from the strain frequency timeseries with

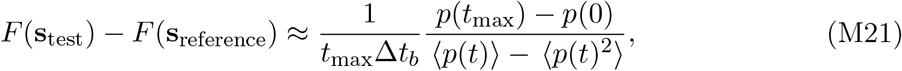

where ⟨·⟩ is a time average. Derivable from a formula by Watterson ^66^, we show how it can be derived in Supplementary Note 5 from the 1-dimensional Kimura equation ^1^, which is the diffusion limit of population genetics.

### 4.8 Iterative fitness landscape design with antibodies (iFLD-A) protocol

The iFLD-A protocol first finds an antibody which binds with high affinity to the starting strain (the WTT antibody). This is accomplished by using simulated annealing as used in the oFLD-A protocol, but with the following main modifications: (1) only a single antibody is used, (2) the antibody is at fixed concentration, so the concentration update steps are irrelevant, and (3) the loss function is set to the binding free energy 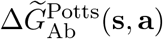. The simulation temperatures are *T*_*MCMC*_ = 0.1 followed by 0.01. The simulated annealing generates the WTT antibody, which we then use to calculate the full fitness landscape over all antigenic sequences. By exhaustively searching the antigenic sequence space of 20^4^ = 160, 000 variants (all amino acid sequences at the 4 antigenic sites) and using the biophysical model in eq. (1) to compute fitness, we identify the peak fitness and its antigenic sequence, which is set as the target sequence, and we once again run simulated annealing to find a high affinity antibody either the globally optimal binder or a deep local optimum for binding affinity. A total of 5 iterations after finding the WTT antibody are performed.

For the cases where we used random starting sequences instead of the true SARS-CoV-2 wildtype, we performed a total of 200 random trials. 20 trials had a starting random (normalized) fitness between 0 and 0.1, 20 trials had starting fitness bewteen 0.1 and 0.2, and so forth. We averaged over all 200 trials to produce Figure 5D.

#### Wright-Fisher viral evolutionary dynamics

Wright-Fisher simulations follow the protocol in Greenbury *et al*. ^5^, where we initialize *N*_pop_ = 1000 antigen sequences at the SARS-CoV-2 wildtype. At each of the 10,000 time steps, we calculate all fitnesses for every individual in the population using eq. (M7). The next generation of 10,000 individuals is selected by sampling the previous generation’s individuals with replacement, with probability 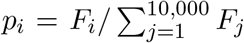, where *i* indexes the previous generation’s individuals. Each of the new generation’s sequence sites are chosen for possible mutation with probability *µ*_chosen_ = 0.01 (for the polymorphic simulations) or *µ*_chosen_ = 0.0001 (for the near-monomorphic simulations). If a site is chosen to mutate, a random amino acid is chosen for that site. This makes the mutation probability *µ* = *µ*_chosen_ ×19/20. The next time step then begins with fitness calculations of all sequences, which are stored. Since for each antigen we consider *L*_Ag_ = 4 residues, the polymorphic dynamics have *N*_pop_*µL*_Ag_ = 38, and the near-monomorphic dynamics have *N*_pop_*µL*_Ag_ = 0.38.

### 4.9 Bounds on population-level fitness

Given our definition of the population-level fitness *F*_pop_(**s**) in eq. (5), the lower bound in eq. (6) was obtained by applying Jensen’s inequality ^78^. An upper bound was obtained by applying Lagrange’s form of Taylor’s remainder theorem ^79^ and then optimizing the bound using the specific form of the *in vivo* biophysical model, eq. (3). The derivation is shown in Supplementary Note 6.

## Acknowledgements

We acknowledge helpful discussions with Anna Sappington, Alexander Kelser, Katherine Ho, Rebecca Greenberg, Aniruddha Chattaraj, Dianzhuo Wang, Bidyut K. Mohanty.

## Funding and resource acknowledgement

This work was supported by a Hertz Foundation Fellowship (VM), and a PD Soros Fellowship (VM), and award T32GM144273 from the National Institute of General Medical Sciences (to Harvard/MIT MD-PhD Program). The content is solely the responsibility of the authors and does not necessarily represent the official views of the National Institute of General Medical Sciences or the National Institutes of Health. We thank the MIT Engaging Cluster, supported by the MIT Office of Research Computing and Data, and the Harvard Faculty of Arts and Science Research Computing (FASRC) for computational resources. The authors declare no known conflict of interest.

## Code and data availability

Code and data will be made publicly available upon publication.

## Supplementary Information

### Supplementary Table

**Table S1:** Mutable antigen and antibody residues used for fitness landscape design.

| Molecule | Mutable Residues |
| --- | --- |
| SARS-CoV-2 Spike Protein | E484, G485, F486, Q493 |
| LY-CoV555 Light Chain | Y32, S91, Y92, S93, T94 |
| LY-CoV555 Heavy Chain | Y101, E102, A103, R104, Y109, Y110 |

### Supplementary Notes

#### Supplementary Note 1: Binding and replication differential equations map onto absolute fitness

##### Binding reactions

We write down the definition of the dissociation constant for eq. (M2):

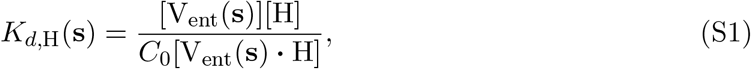

and for eq. (M3)

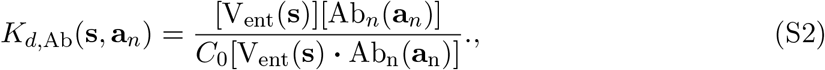

where *C*_0_ is the reference concentration used to make the dissociation constants dimensionless. We seek an expression for *p*_*b*_(**s**), the equilibirum host-bound fraction of viral entry proteins:

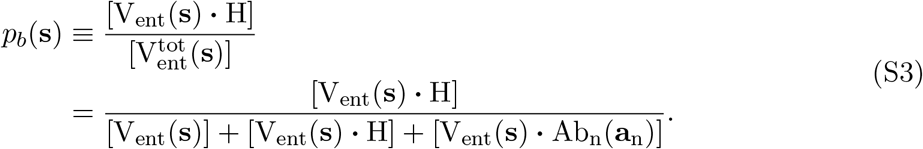

Substituting using eq. (S1) and eq. (S2), we obtain:

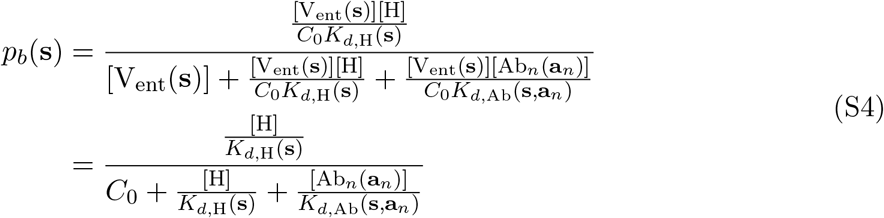

Using Δ*G*_H_(**s**) = *β*^−1^ log *K*_*d*,H_(**s**) as well as Δ*G*_Ab_(**s, a**_*n*_) = *β*^−1^ log *K*_*d*,Ab_(**s, a**_*n*_), we arrive at

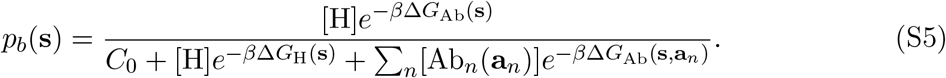

[H] and [Ab_*n*_(**a**_*n*_)] are unbound host and antibody concentrations.

We assume that the availability of infectable host cells and receptors is much higher than the viral load, and clinical data suggests that antibody concentrations are much higher than the viral concentration in a immunocompetent, vaccinated individual (Supplementary Note 2). Assuming that host receptors and antibodies are not significantly depleted by viral binding, we can approximately equate the free and experimentally controllable total concentrations [H^tot^] ≈ [H] and 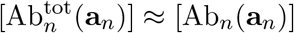. Thus, we have

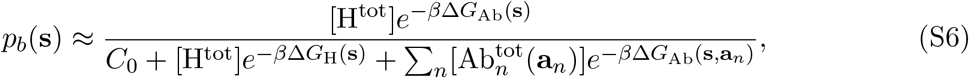

which is eq. (2).

##### Replication dynamics

We note that the chemical reaction eq. (S7) produces *unbound* copies of the virus, which we temporarily denote as *V* (**s**)·(0 H) for mathematical convenience. Thus, we have

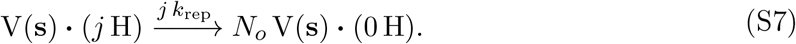

The ordinary differential equation for the time evolution of the unbound concentration must include contributions from the rate equations for values of all *j* ∈ {1, …, *N*_ent_}:

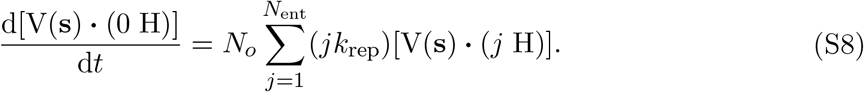

The same reaction equation also provides the depletion rate of the replicating virus bound to *j* host receptors:

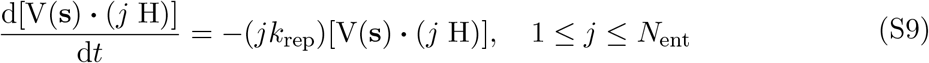

We now denote the total viral concentration for strain **s** as

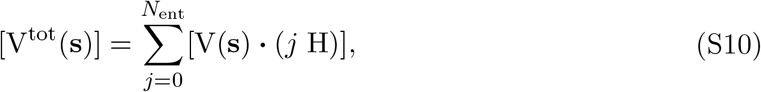

which includes all viruses in all bound and unbound states. Taking a time derivative, it immediately follows that

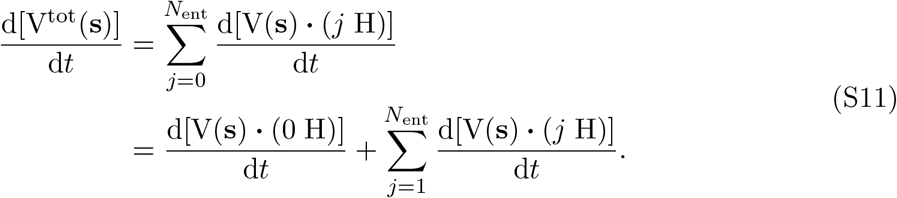

Plugging in the results of eq. (S8) and eq. (S9), we immediately have

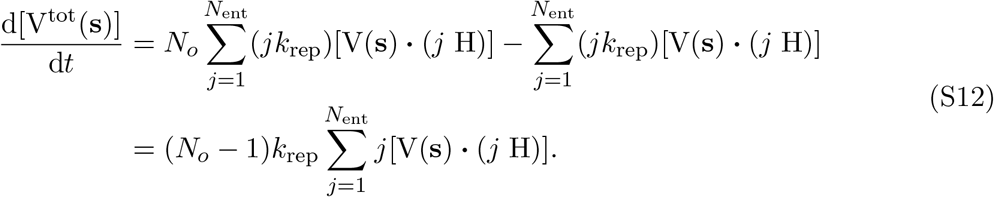

We now invoke separation of timescales: if the binding dynamics occur at timescales much faster than replication, then [V(**s**) · (*j* H)] can be calculated at equilibrium. At equilibrium, we showed that the probability of finding any viral entry protein bound to a host receptor is *p*_*b*_(**s**). Thus, the number of entry proteins on a single virion which are bound to host receptors is a binomial random variable *J* ~ Binomial(*N*_ent_, *p*_*b*_(**s**)). The fraction of virions bound to *j* hosts should follow the distribution of *J*:

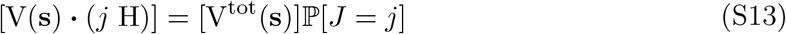

Substituting into eq. (S12), it follows that

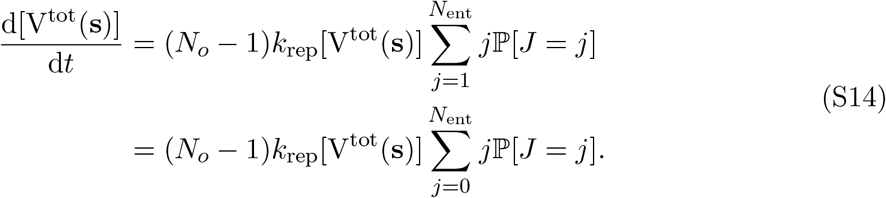

Noting that the sum is exactly the expectation of a binomial variable, we have

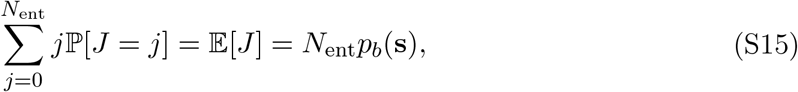

which provides the final result

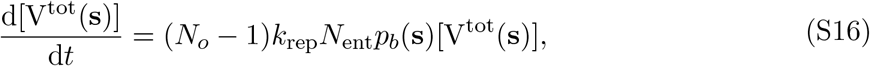

which is eq. (M5), the desired result.

#### Supplementary Note 2: Justficiation of antibody and viral receptor relative concentrations from clinical data

We note that a 50-year-old’s mean peak viral load from a SARS-CoV-2 infection—independent of vaccination status—is 10^8.14^ ≈ 1.28×10^8^ copies/mL ^80^. Each SARS-CoV-2 virion has, on average, 24 *±* 9 spike protein trimers, so there are on average 72 = 7.2 × 10^1^ spike proteins per virion ^81^. Together, this suggests that an infected individual has [V_ent_(**s**)] ≈ (1.28 × 10^8^) × (7.2 × 10^1^) ≈ 9.22 × 10^9^ proteins/mL.

Recent clinical data ^82^ shows that previously uninfected individuals who received 2 doses of the SARS-CoV-2 mRNA vaccine had median spike protein-specific IgG concentration of 46.80 *µ*g/mL, with previously infected but unvaccinated invidiuals having roughly 1 order of magnitude lower concentration and vaccinated individuals who had also been previously infected having roughly 1 order of magnitude greater concentration. Since IgGs have an average molecular weight of 150 kDa (kg/mol) ^83^, we approximately have

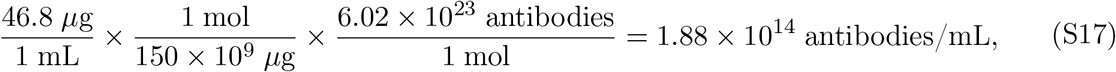

suggesting that antibody concentration outweighs viral concentration. Even with up to 10^2^-10^3^ orders of magnitude of epitope diversity, the estimated antibody concentration outweighs SARS-CoV-2 spike proteins.

The total host concentration is the sum of bound and unbound host concentrations [H^tot^] = [H] + [V_ent_(**s**) · H], and the total antibody concentration is the sum of bound and un-bound host concentrations 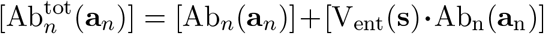. Since we estimate antibody concentrations are greater than the number of viral proteins by orders of magnitude and assume vast availability of infectable host cells, we can write 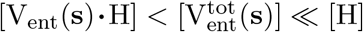 and 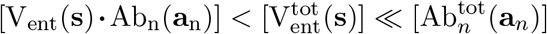. Therefore, we can approximately equate the free and experimentally controllable total concentrations [H^tot^] ≈ [H] and 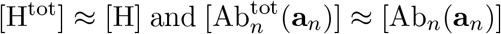.

#### Supplementary Note 3: In vivo biophysical fitness model, with immune-mediated clearance and multiple epitopes, from chemical reactions

We now extend the biophysical model in Supplementary Note 1 to include multiple binding sites on the viral entry/fusion protein as well as reactions for immune-mediated clearance of virions.

##### Binding reactions

Suppose that every antigen has *N*_*S*_ unique binding sites, and we suppose that the *i*-th site can be bound by 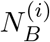 unique chemical species, given by the set 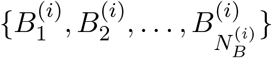. Without loss of generality, we designate site *i* = 1 to be the site where the host receptor binds, and we simply assign a convention 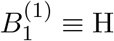 to aid with labeling.

We write down the definition of the dissociation constant for entry protein site *i* and some binding species 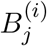:

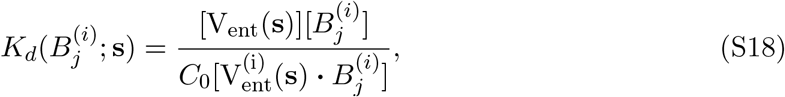

where, again, *C*_0_ is the reference concentration used to make the dissociation constants dimensionless. We seek an expression for 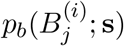, the fraction of viral entry proteins whose *i*-th site is bound to species 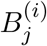 at equilibrium:

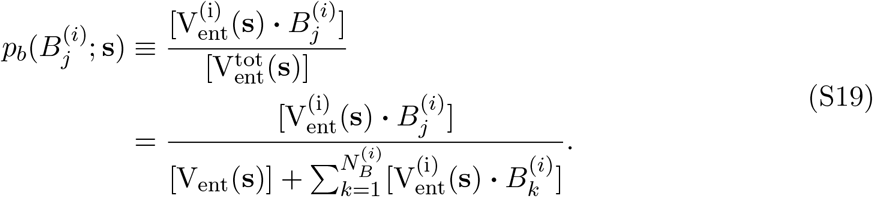

Substituting using eq. (S18), we obtain

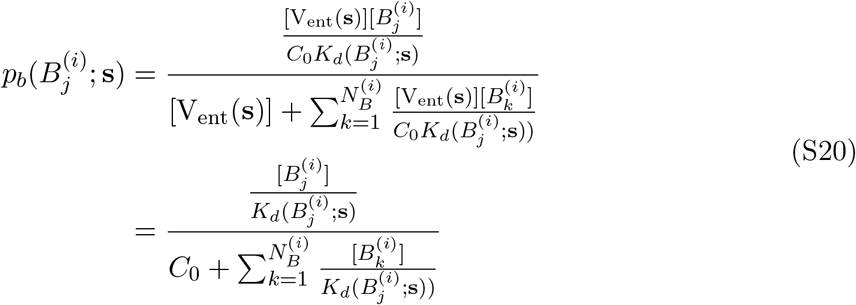

Using 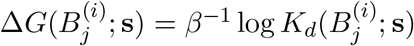, we arrive at

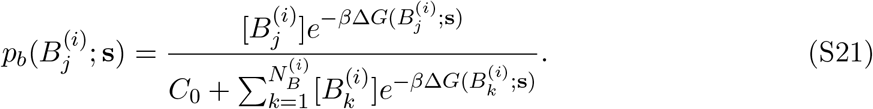

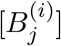 is the unbound concentration of 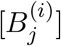. This is the standard Langmuir adsorption isotherm also derivable from standard adsorption models in statistical mechanics. Once again, working under the approximation that the total amount of chemical binders (e.g. hosts and antibodies) greatly exceeds the number of virions, as assumed previously in Supplementary Note 1 and rationalized in Supplementary Note 2, we have 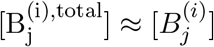, which gives us

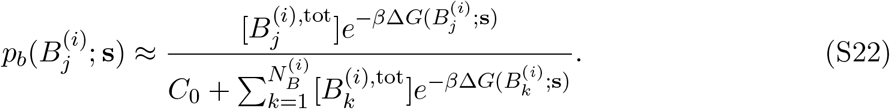

The probability of the site being unbound is accordingly

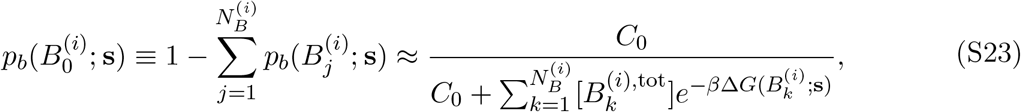

where we defined 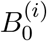 to be the state where *i* is not bound.

Next, under the assumption that chemical binders are site-specific (so an antibody which binds to site 1 will not bind to site 2, for instance), every site would equilibrate independently. Thus, it immediately follows that the probability that a single entry protein has some binding state given by 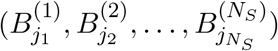, with 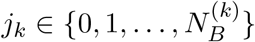, is the product of each site’s binding probabilities

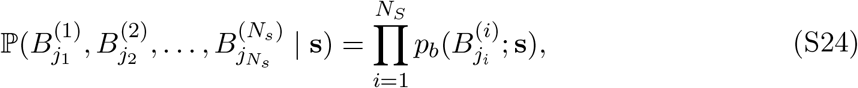

where *j*_*i*_ represents which species is bound to site *i* (or potentially the state where no species is bound).

Now, consider a single complete virion, which has *N*_ent_ entry/fusion proteins, which we assume equilibrate independently of each other. Let 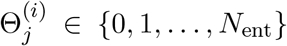 represent the total *number* of proteins whose site *i* is bound to species 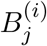, again letting *j* = 0 represent the unbound state. Then, 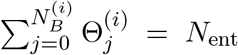 because every site on every protein either must be bound to one of the 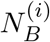 chemical species or must be unbound. For any site *i*, the probability mass function for the binding states across all entry proteins is given by a multinomial distribution

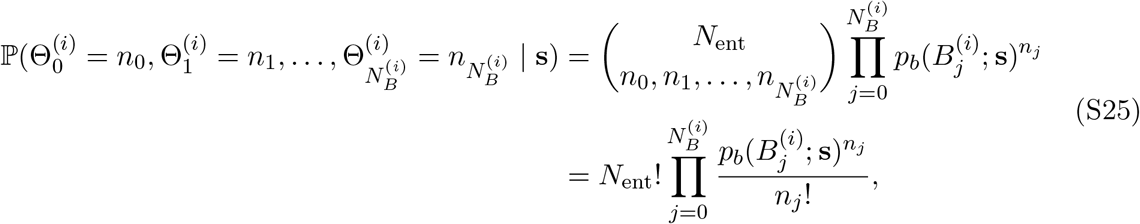

With 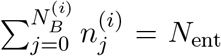. Extending this to all sites, which equilibrate independently of each other, the joint probability mass function for the binding states across all entry proteins is a product of these multinomial distributions

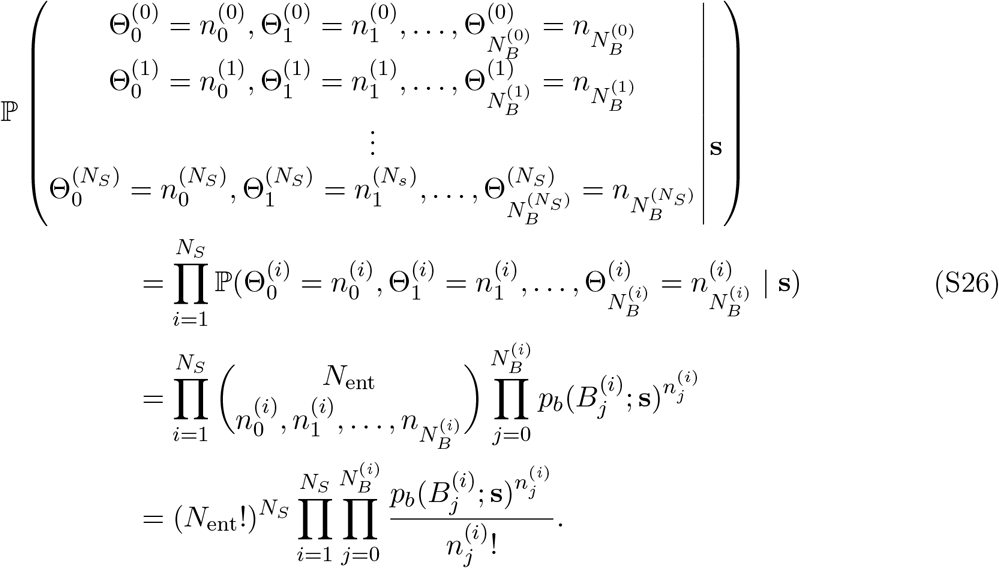

We now consider the form of a marginal distribution of a single 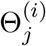 (i.e. the distribution of the number of 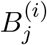 molecules which are bound to a single virion). This is obtained by summing over all binding number configurations 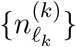 over all sites *k*≠ *i*, and by summing over all binding number configurations for species *not* of type 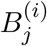 at site *i*:

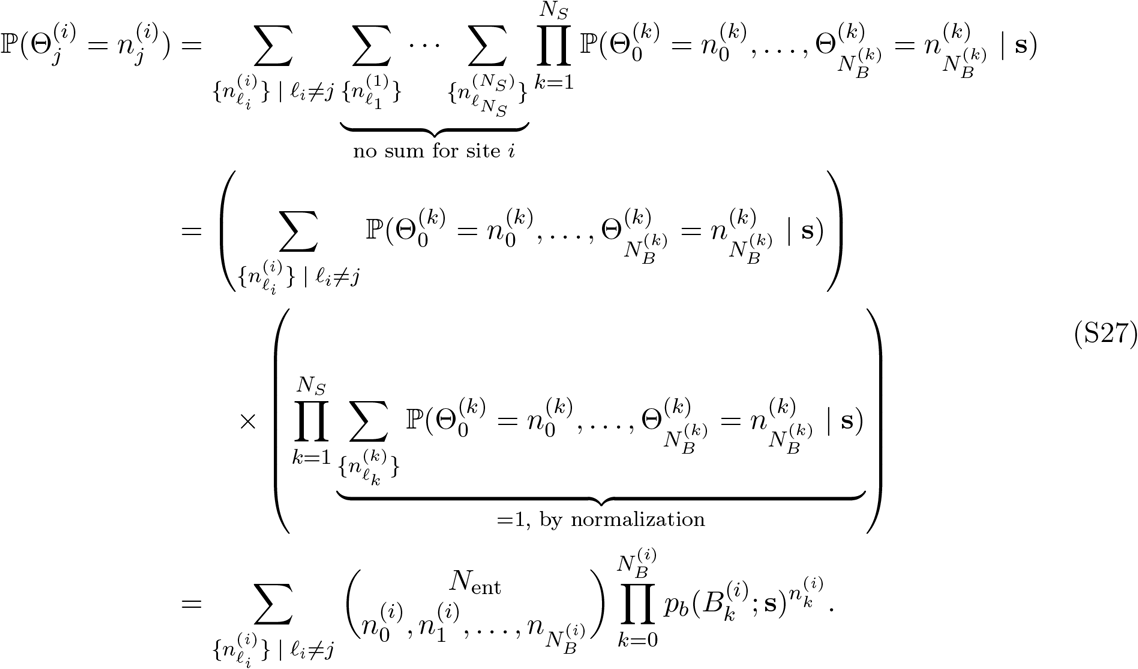

Now, we will show that marginalizing over all possible values of 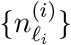 for *ℓ*_*i*_≠ *j* will turn the multinomial distribution into a binomial distribution. Starting by expanding the multinomial coefficient above, we first bring the *j*-dependent terms outside the summation

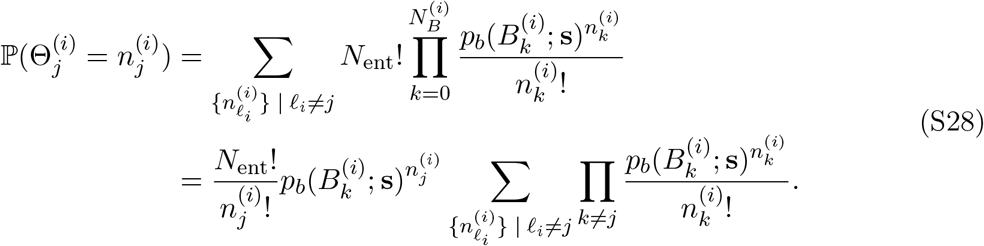

Note that from the multinomial theorem, we would have

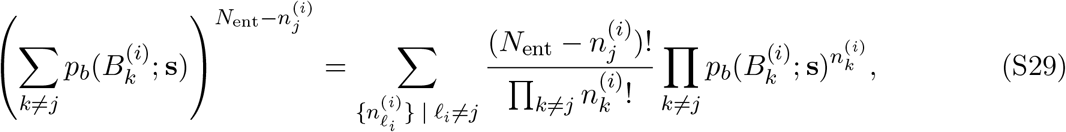

which can be rearranged to

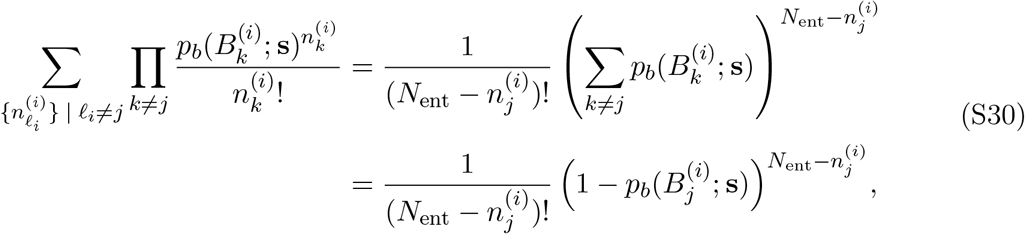

where in the last line we have used the fact that

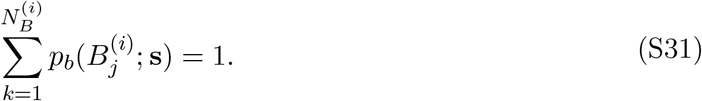

It now follows that

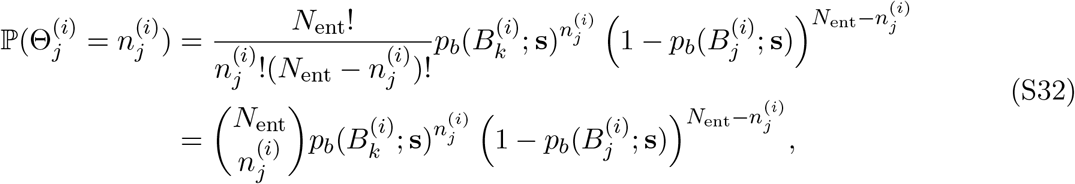

which is exactly a binomial distribution. Therefore, the marginal probability that 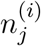 of the *N*_ent_ entry proteins’ sites of type *i* are bound by species 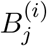 is given by a binomial distribution which depends on the equilibrium probability 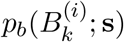 of that species being bound, which was calculated earlier in this section. Recall that the expectation of the binomial distribution is given by the simple formula

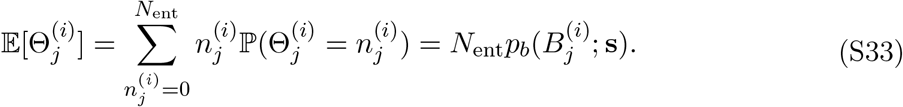

This applies regardless of the type of species considered, whether host receptor or antibody; the above result will be useful in the following section where the biophysical fitness model is derived.

##### Replication dynamics

We now seek to derive an expression for fitness by directly calculating the growth rate 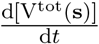 of the total viral population for a strain **s**. Categorizing the viral population based on the binding state configurations 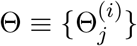 defined in the previous section, we can write

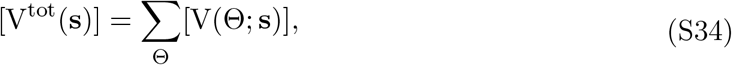

so

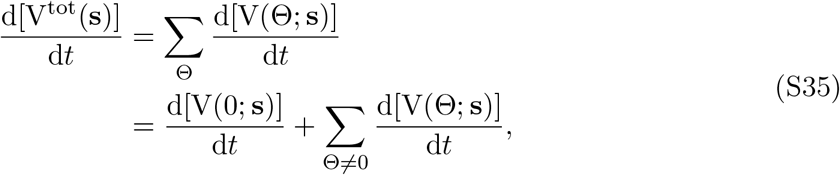

where Θ = 0 indicates virions which are completely unbound. As done in Supplementary Note 1, we take all new viral offspring to be completely unbound, which means that 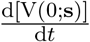 is impacted only by the chemical reaction for replication. It also cannot be cleared since antibodies are not bound. Recall the convention from the previous section that species 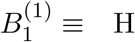 represents the host receptor, and the host receptor only binds viral protein site 1. From our chemical reaction model, we assume that if *θ* host receptors are bound, the rate constant for the replication reaction in which *N*_*O*_ offspring are produced is *θk*_rep_. Thus, we have

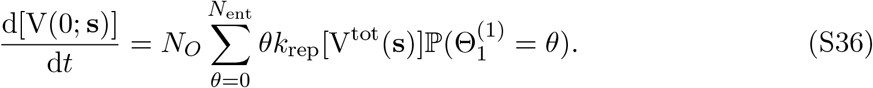

Here, we have assumed that the concentration of viruses bound to *θ* hosts can simply be expressed as the total viral concentration [V^tot^(**s**)] times the probability of a virion being bound to *θ* hosts 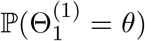. Using the eq. (S33) derived previously, we have

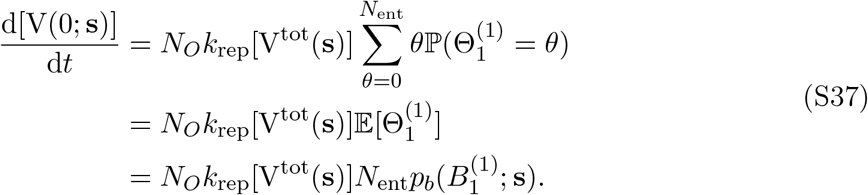

Now, we consider Θ≠ 0, so at least one species is bound. These could be host receptors, antibodies, or both. We note that from the replication chemical reactions in the Methods section, if *θ* host receptors are bound, the rate constant for the replication reaction in which one virion is destroyed during the replication process is *θk*_rep_. Also, from the clearance chemical reactions in the Methods section, if *φ* antibodies are bound (of any kind, at any site), the rate constant for the clearance reaction in which one virion is cleared during the replication process is *φk*_cle_. Clearance must be considered for all antibodies at all sites as well. We consider site 1 separately from other sites, since 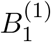 is the host receptor. Thus, we can now write, for Θ≠ 0,

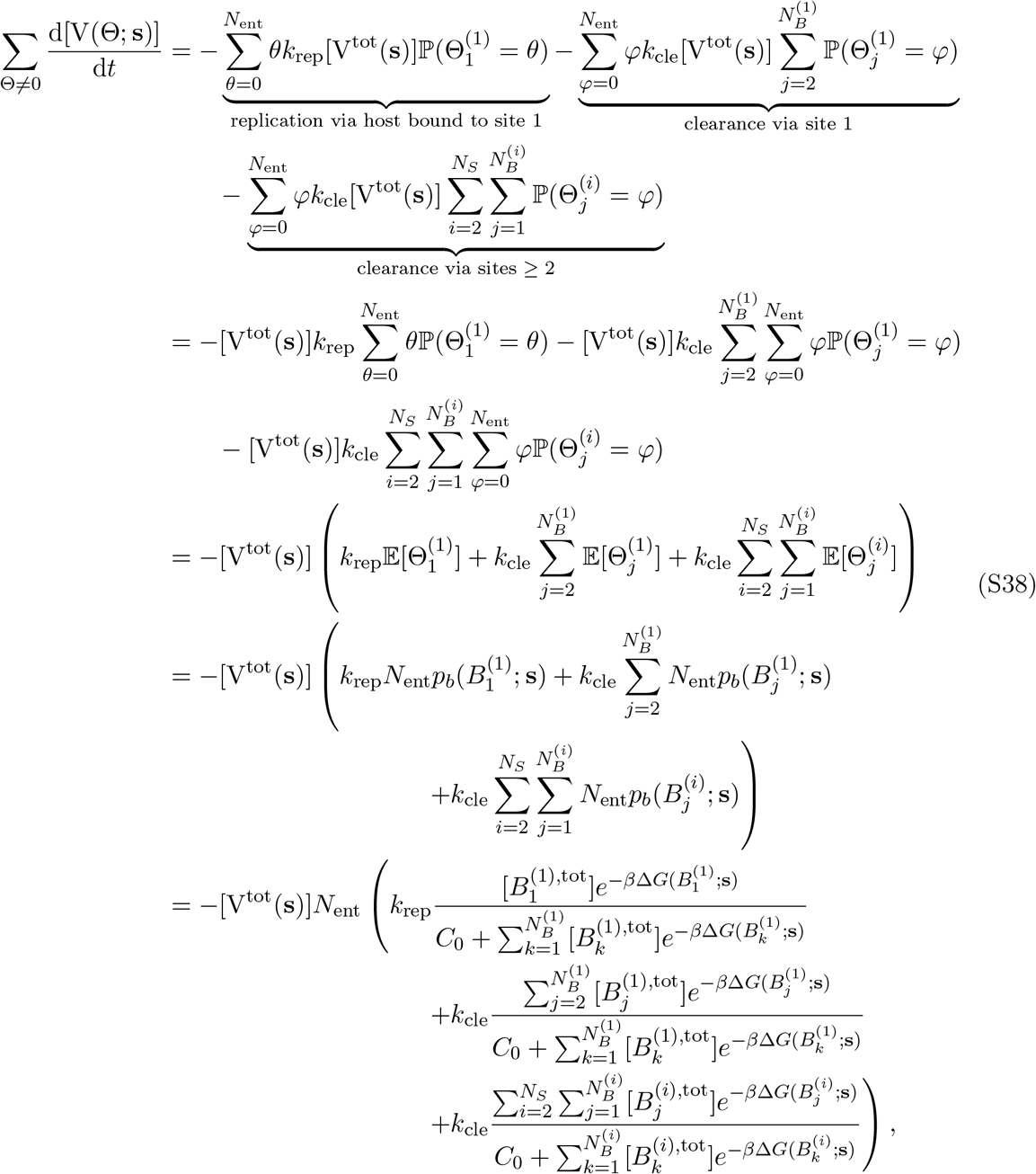

where in the first line the sums over *θ* and *φ* can start at zero despite Θ≠ 0 because the summand is zero when *θ* = 0 or *φ* = 0. We now replace the general notation with host and antibody labels: 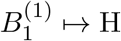 represents the host receptors, 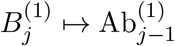 represents antibodies which directly compete with the host for binding to site 1 (with numbering scheme adjusted), and 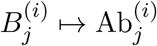 represents antibodies which do not directly compete with the host receptor for binding to the antigen (and bind to other epitopes *i*≠ 1). Thus, we can write

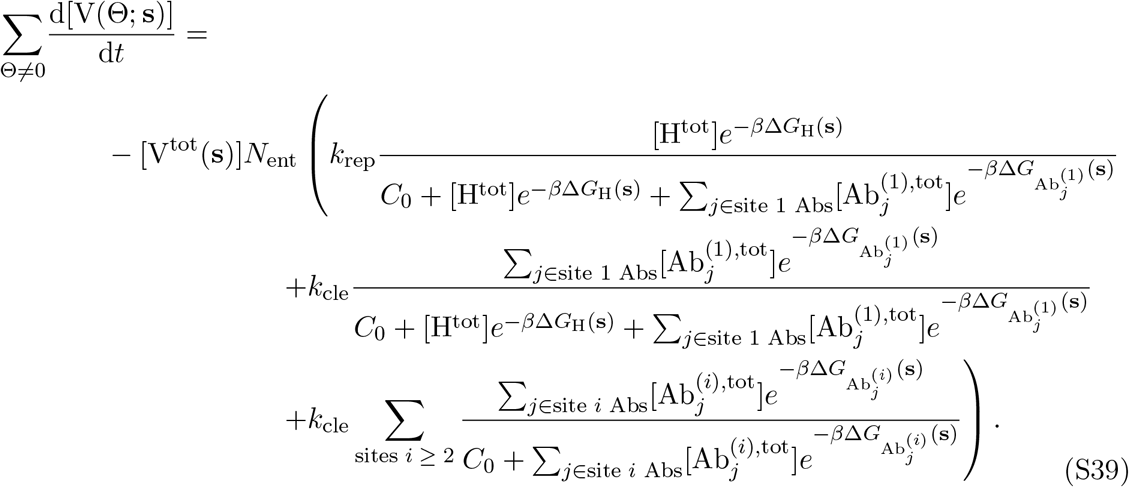

Substituting eq. (S37) and eq. (S39) into eq. (S35), we have

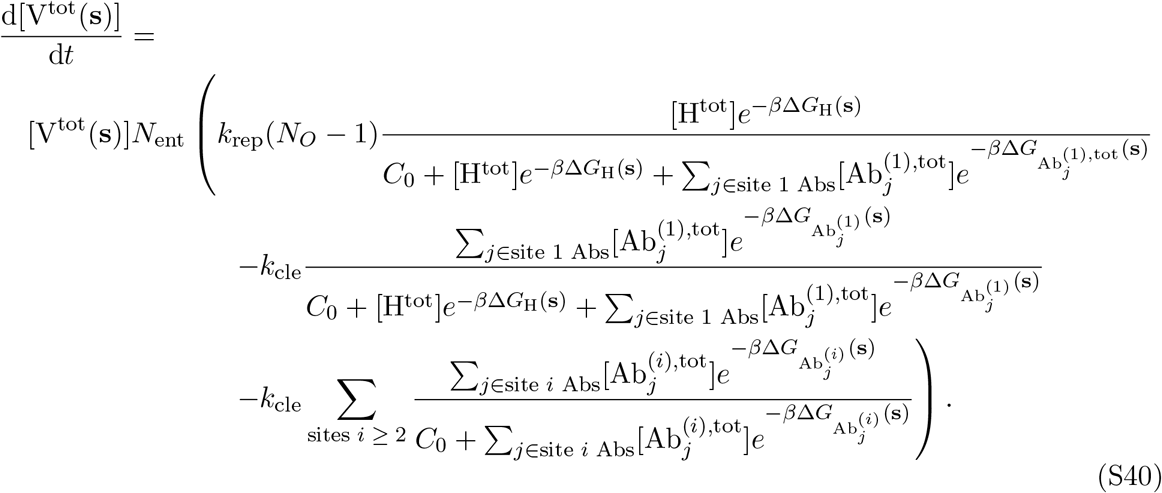

Again, using the definition of absolute fitness

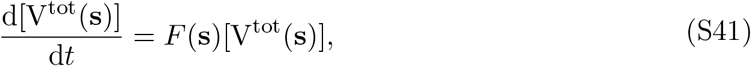

we can now write the fitness, now including immune-mediated clearance as well as multiple antigenic epitopes, as an extended version of the *in vitro* biophysical fitness derived in Supplementary Note 1:

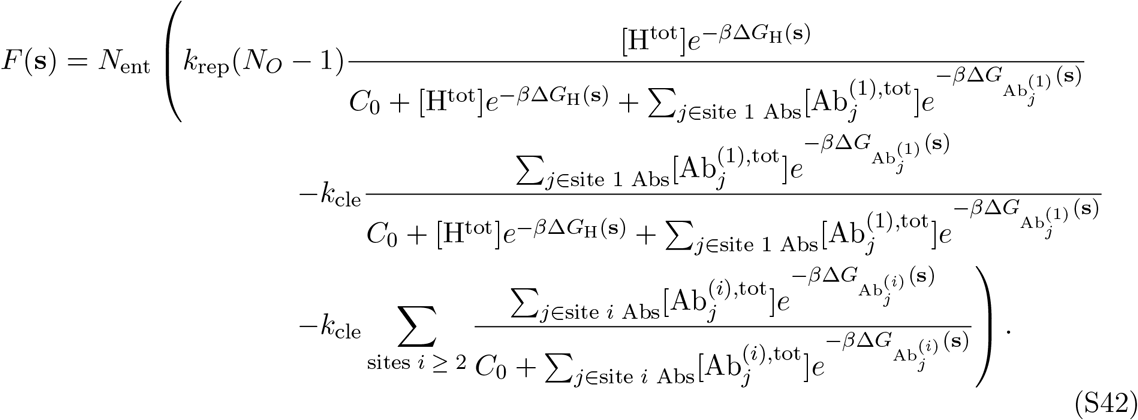

#### Supplementary Note 4: Relation to hypothesized biophysical model (Wang et al., **Proc. Natl. Acad. Sci**., **2024)** ^**18**^

We now show how the above *in vivo* model, which includes immune clearance, relates to our previously hypothesized biophysical model ^18^, which included modeling of SARS-CoV-2 spike receptor binding domain up/down configurations. This model was fit to fitness labels obtained from Obermeyer *et al*. ^60^ and obtained high accuracy. It had been shown that spike receptor binding domain up/down configurations do not affect the analytical form of the model, which took the form

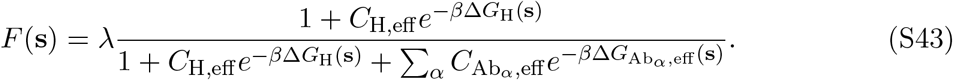

Here, *C*_H,eff_ is an effective renormalized concentration of the host receptor in the population, 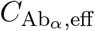 is the effective renormalized concentration of antibody *α* in the population, and Gibbs free energies represent the binding free energy to the receptor binding domain “up” configuration. The model had been hypothesized in this form, but it has never been shown how it may arise directly from non-equilibrium chemical reaction dynamics.

Now, we note that in our explicitly derived biophysical model, suppose we take all antibodies to compete directly with the host receptor for binding. This means that a spike protein’s receptor binding domain bound to an antibody cannot bind to a host receptor, perhaps due to steric clash. Then, eq. (S42) becomes (dropping the site 1 index):

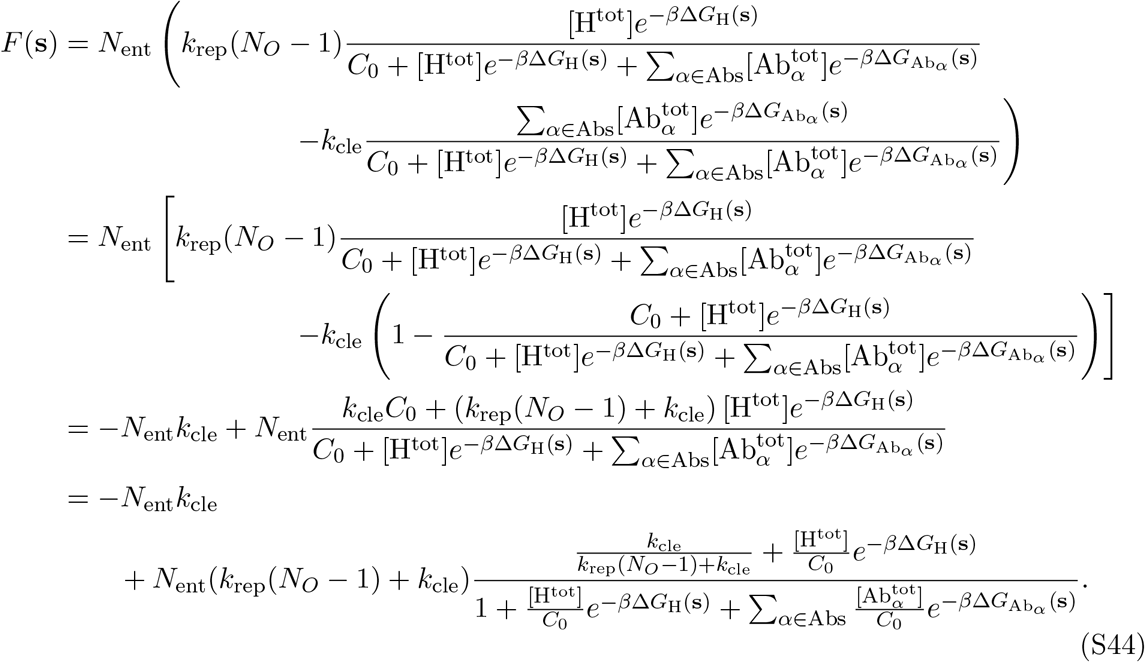

Malthusian *relative* fitnesses impact evolutionary dynamics, so constant shifts of absolute fitness *F*(**s**) ↦ *F*(**s**) + *A* are not expected to lead to changes in relative abundances of genotypes. Thus, defining a relative fitness *F*_rel_(**s**) = *F*(**s**) + *N*_ent_*k*_rep_ and setting 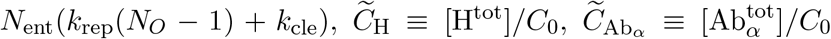, and 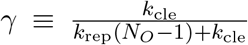, we have

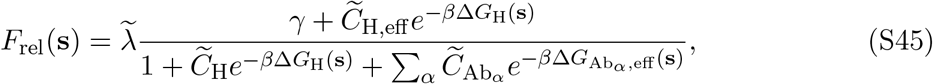

and 0 < *γ* < 1 for nonzero *k*_rep_ and *k*_cle_.

It is easy to see that in the limit of *γ* = 0 (weak clearance limit, *k*_cle_ ≪ *k*_rep_(*N*_*O*_ − 1)), the *in vitro* model from Supplementary Note 1 is exactly recovered, as would be expected from the fitness model eq. (S42). In the limit of *γ* = 1 (strong clearance limit, *k*_cle_ ≫ *k*_rep_(*N*_*O*_ − 1)), the hypothesized biophysical model from our previous work ^18^, eq. (S43), is exactly recovered. In the intermediate case where 0 < *γ* < 1, but *γ* is not near one of its extremal values, we essentially have a model which is nearly identical to eq. (S43), but contains an extra free parameter, which would naturally cause this model to fit the PCR-derived SARS-CoV-2 fitness data better than, or equally as well as, the model from our previous paper ^18^. The implications are discussed in the main text.

#### Supplementary Note 5: Fitness estimation from strain frequency time series

When there are two strains (test and reference), and *p*(*t*) is the test strain frequency as a function of time *t*, the mean rate of change of the strain frequency—in continuous time— is given by the 1-dimensional Kimura equation ^1^ in the limit of no stochastic fluctuations (equivalently, the infinite population limit):

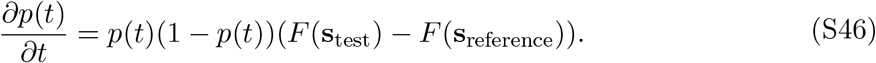

First, we account for the fact that the *in silico* serial dilution experiment timescale is different from the broth dilution time scale. Any exponential growth during a time Δ*t*_*b*_ in the simulation time looks like unit time in the broth time series. Thus, the fitnesses in the equation above need to be multiplied by a factor of Δ*t*_*b*_. Thus, we have

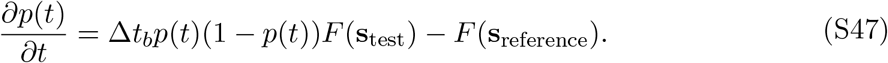

Now, time averaging over both sides of the equation from [0, *t*_max_], we have

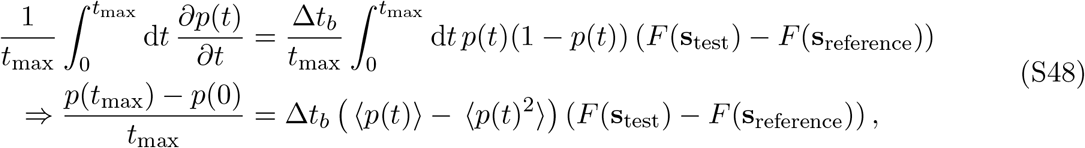

which is rearranged to

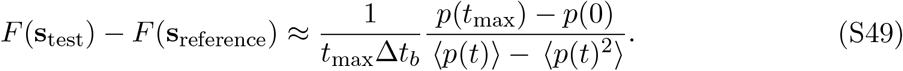

This can be derived from a formula by Watterson ^66^. The formula derived with forward-time finite difference will give time averages which are off by a boundary term in the series. We have not investigated the effect of deriving the above formula with an integral versus a sum but do not expect fitness estimation to be substantially affected.

#### Supplementary Note 6: Proof of bounds on population-level fitness

Starting with Main Text eq. (5), we can collapse the sum over vaccine antigens to get an expectation that reads as an integral over the antibody concentrations

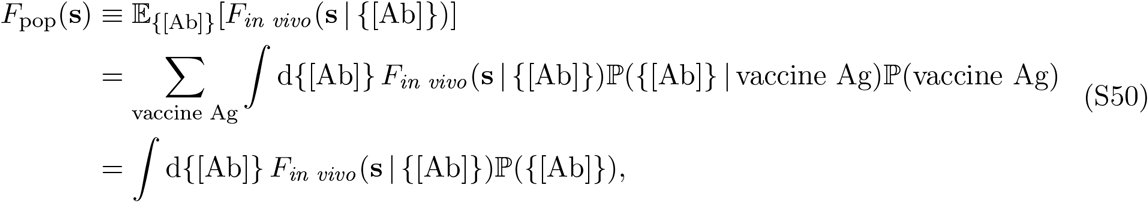

where *F*_*in vivo*_(**s** | {[Ab]}) is the *in vivo* biophysical model from Main Text eq. (3), reproduced below

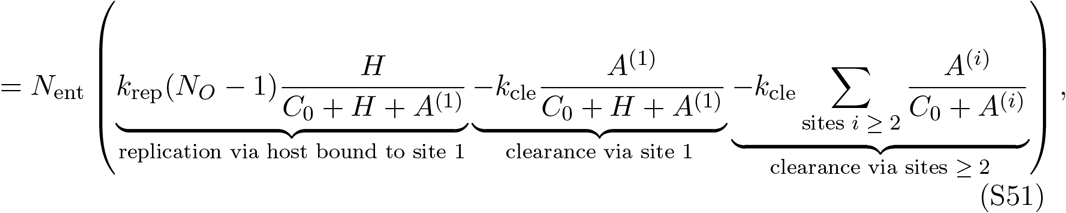

with

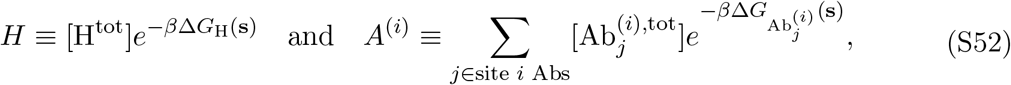

We define some more compact notation for convenience: let 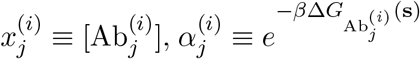, *C*_*H*_ ≡ *C*_0_ + *H, k*_*R*_ ≡ *N*_ent_*k*_rep_(*N*_*O*_ − 1), *k*_*C*_ ≡ *N*_ent_*k*_cle_, *F* ≡ *F*_*in vivo*_(**s** | {[Ab]}),

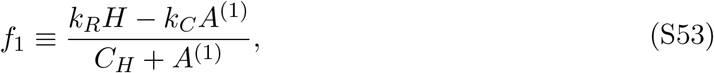

and

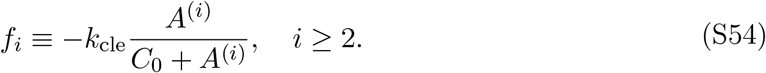

Now, we have

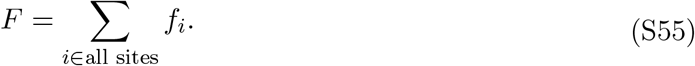

It is assumed that the host concentration is constant across the population, but antibody concentrations are the random variables whose joint distribution is ℙ ({[Ab]}) = ℙ (**x**).

We now assess the curvature (i.e. Hessian) of the fitness function with respect to any two antibody concentrations. First, consider two sites *k* and *ℓ*. The derivative with respect to a site *k* antibody concentration

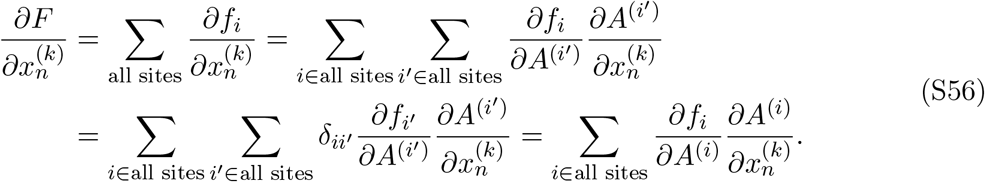

Noting that 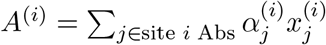, we have

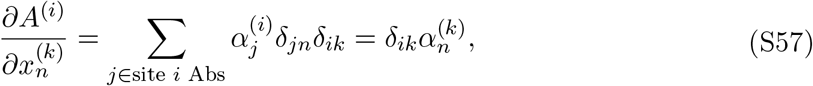

which gives us

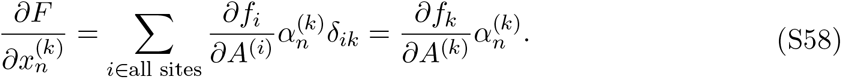

Taking another derivative with respect to antibody concentration, we have

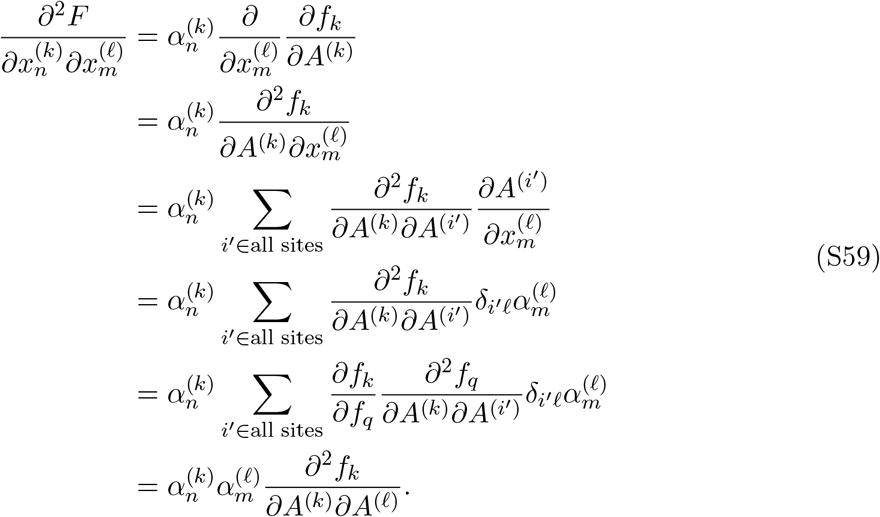

Note that since each

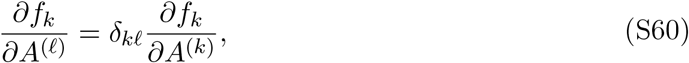

we can write

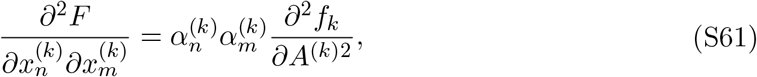

and

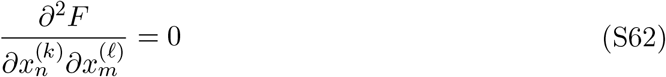

for *k*≠ *ℓ*. Thus, the Hessian matrix has block diagonal structure, with each block corresponding to an epitope/binding site.

Now, we can write

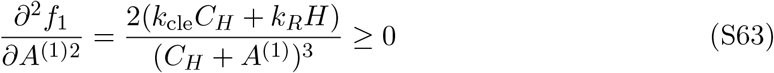

and

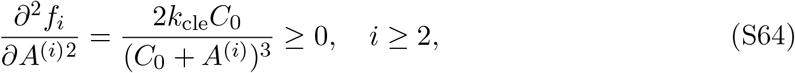

which means that the Hessian is positive semi-definite. Thus, the *in vivo* fitness function is globally convex over the domain of all antibody concentrations. This means that Jensen’s inequality^78^ provides a lower bound on the population-level fitness

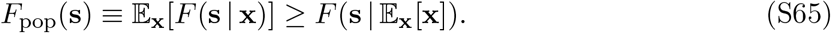

To establish an upper bound, we Taylor expand the *in vivo* fitness around the population-averaged antibody concentrations 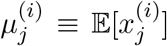 and using Lagrange’s representation of the remainder ^79^,

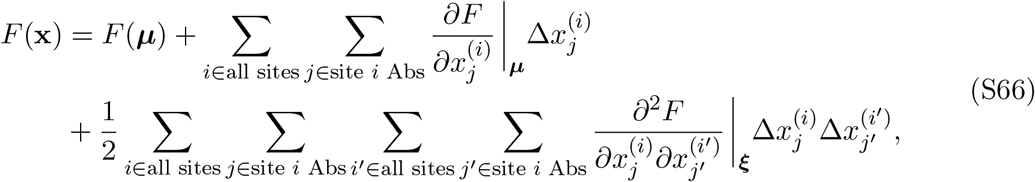

where we have defined 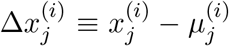, and ***ξ*** = ***µ*** + *t*(**x** − ***µ***) for *t* ∈ (0, 1) is some point collinear with **x** and ***µ*** and is collinear with these two points. Taking the expectation of both sides, we have

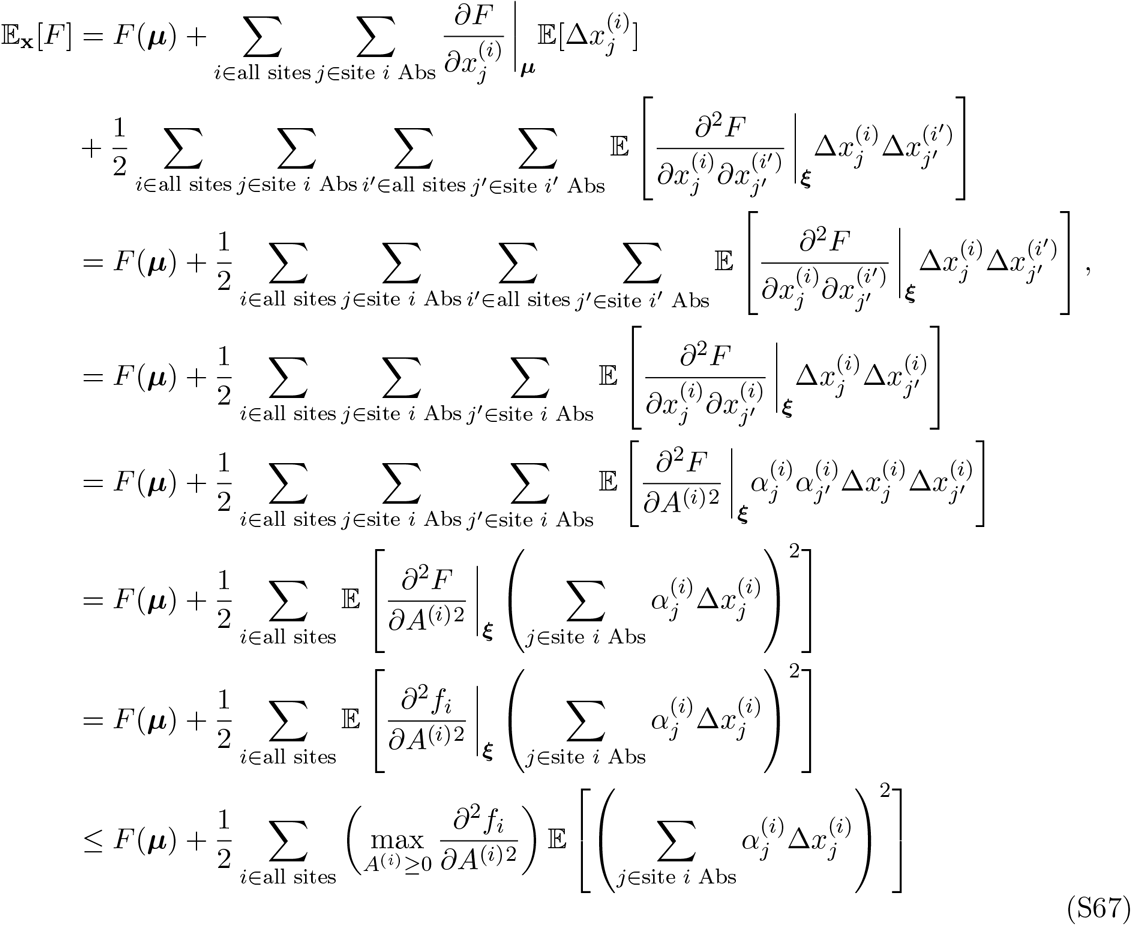

where in the first line we have used the fact that the first-order term vanishes since 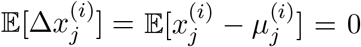, and. In the last line we have noted that the partial derivative evaluated at ***ξ*** is maximized when the antibody concentrations are zero anyway: from eq. (S63) and eq. (S64), we see that the second derivatives with form ~ 1/(*C*_0_ + *A*^(*i*)^)^3^ are maximized when antibodies are absent *A*^(*i*)^ = 0:

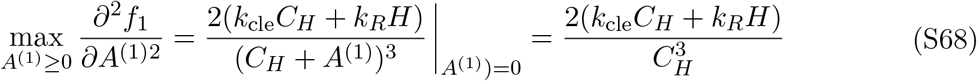

and

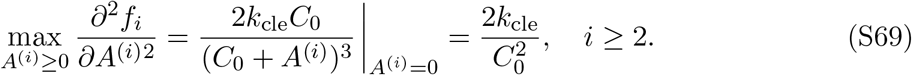

Also, we define the Boltzmann-weighed variance of the antibody concentration at site 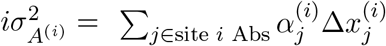. Putting these results, together, we have

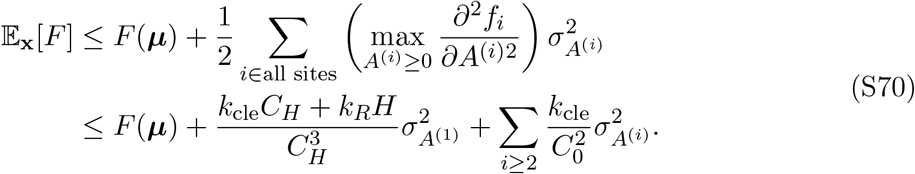

This provides an upper bound on the population-level fitness. We can now write

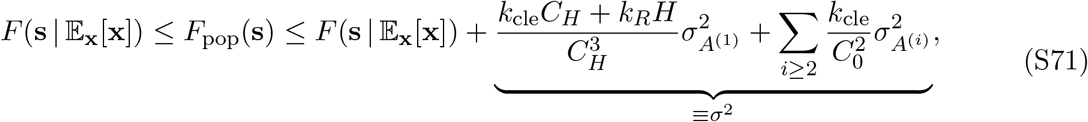

where *σ*^2^ is the weighted variance-like term indicated in Main Text eq. (6). This completes the proof.

## Supplementary Figures

**Figure S1:**
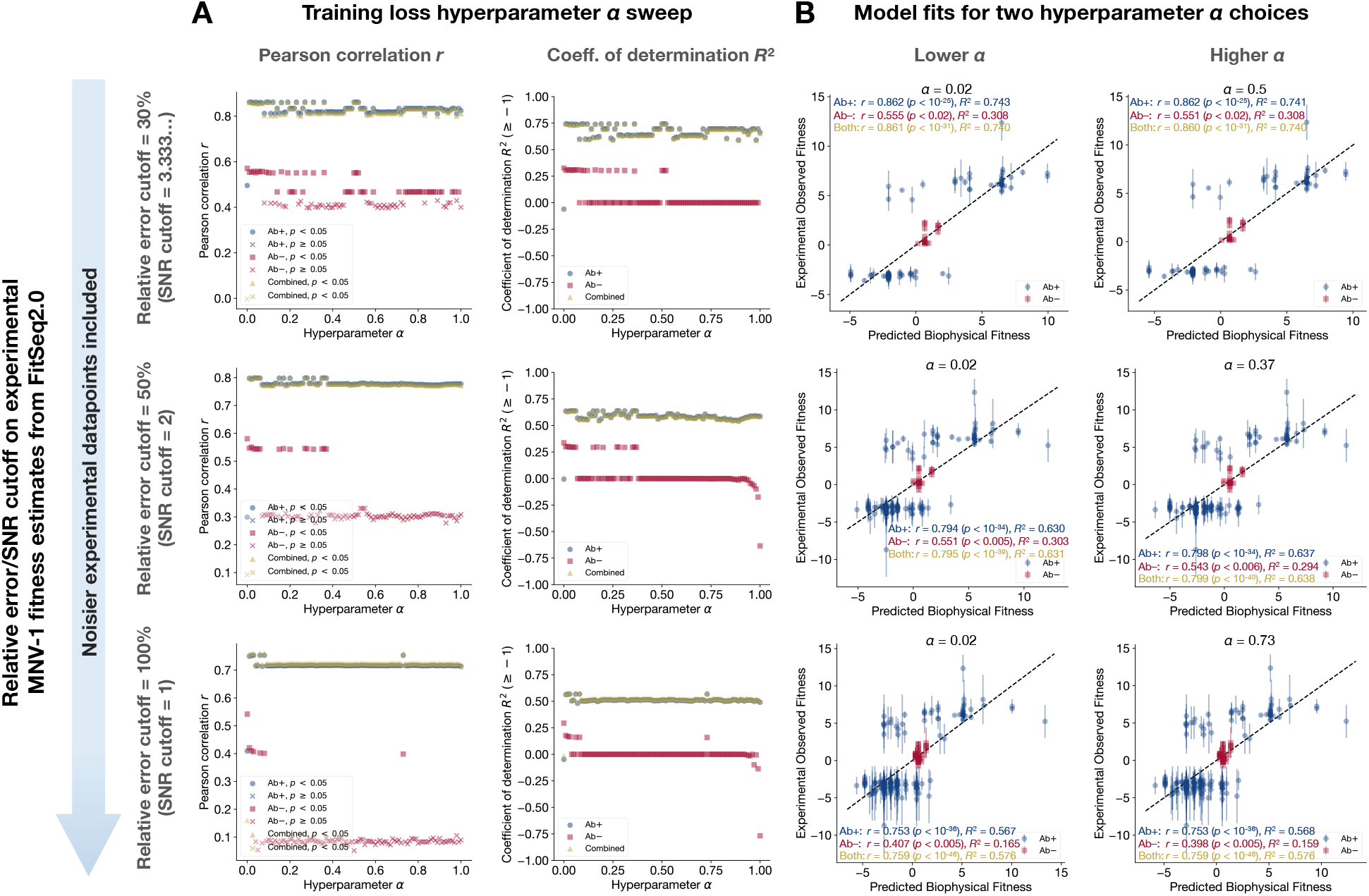
Validation of *in vitro* biophysical model (*with* free scale parameter) on experimental fitnesses from MNV-1 serial passage experiments using EvoEF-predicted binding free energies. **(A)** We show Pearson correlation *r* and coefficient of determination *R*^2^ plotted against training hyperparameter *α*, where the goodness-of-fit measures are for fitting the biophysical model to FitSeq2.0-derived experimental fitness measurements from serial passage experiments with and without antibodies, and for the combined dataset (**B**) For two different choices of training hyperparmaeter *α*, we show FitSeq2.0-derived experimental fitness measurements plotted against biophysical model fitness, where biophysical fitness calculations use EvoEF-derived free energies. For both (A) and (B), different relative error/SNR cutoffs on the FitSeq2.0-derived experimental fitness measurements are shown in each row. The top left panel of (B) is reproduced in the Main Text as Figure 3A.

**Figure S2:**
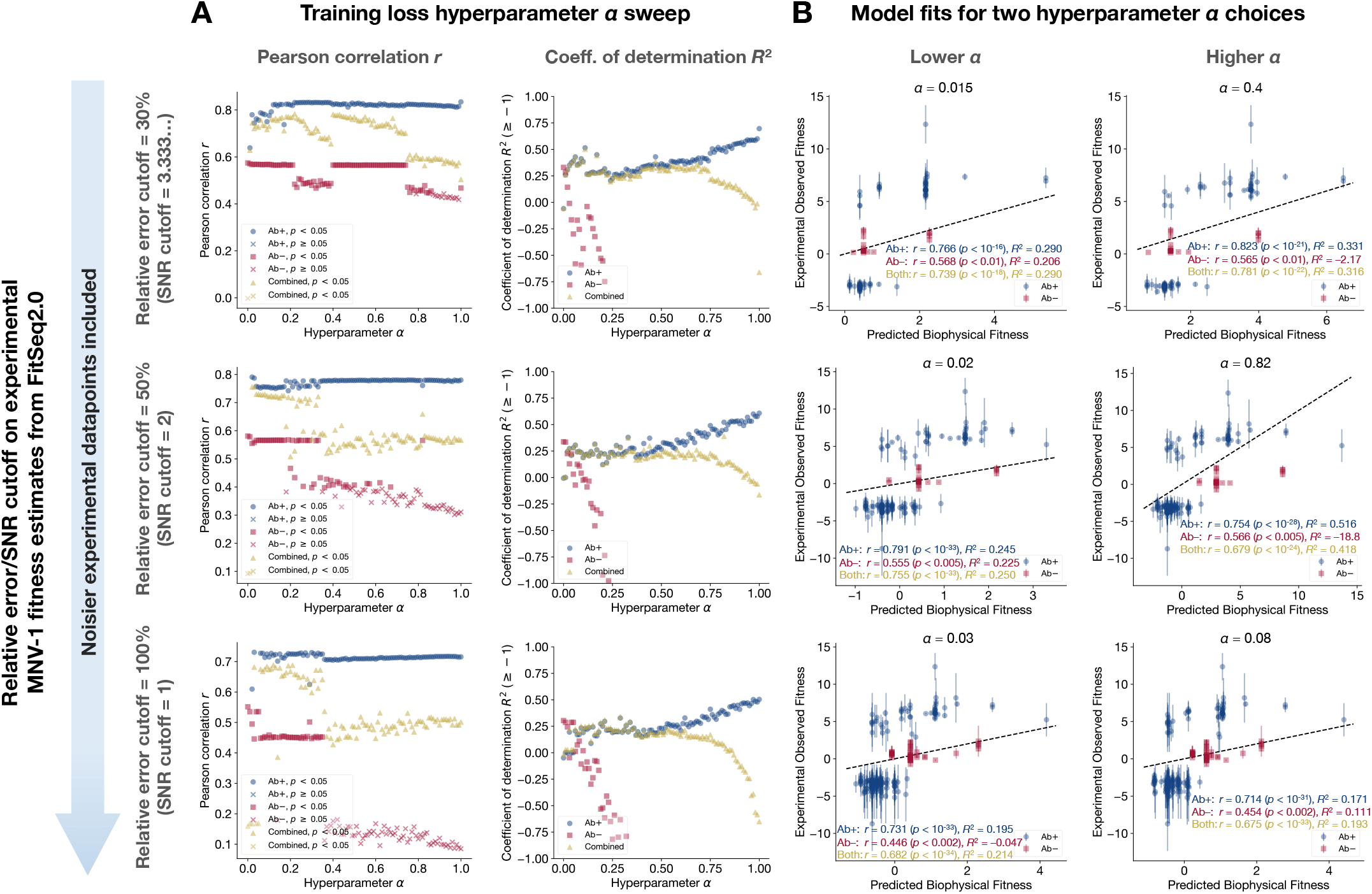
Validation of *in vitro* biophysical model (*without* free scale parameter) on experimental fitnesses from MNV-1 serial passage experiments using EvoEF-predicted binding free energies. (**A**) We show Pearson correlation *r* and coefficient of determination *R*^2^ plotted against training hyperparameter *α*, where the goodness-of-fit measures are for fitting the biophysical model to FitSeq2.0-derived experimental fitness measurements from serial passage experiments with and without antibodies, and for the combined dataset (**B**) For two different choices of training hyperparmaeter *α*, we show FitSeq2.0-derived experimental fitness measurements plotted against biophysical model fitness, where biophysical fitness calculations use EvoEF-derived free energies. For both (A) and (B), different relative error/SNR cutoffs on the FitSeq2.0-derived experimental fitness measurements are shown in each row.

**Figure S3:**
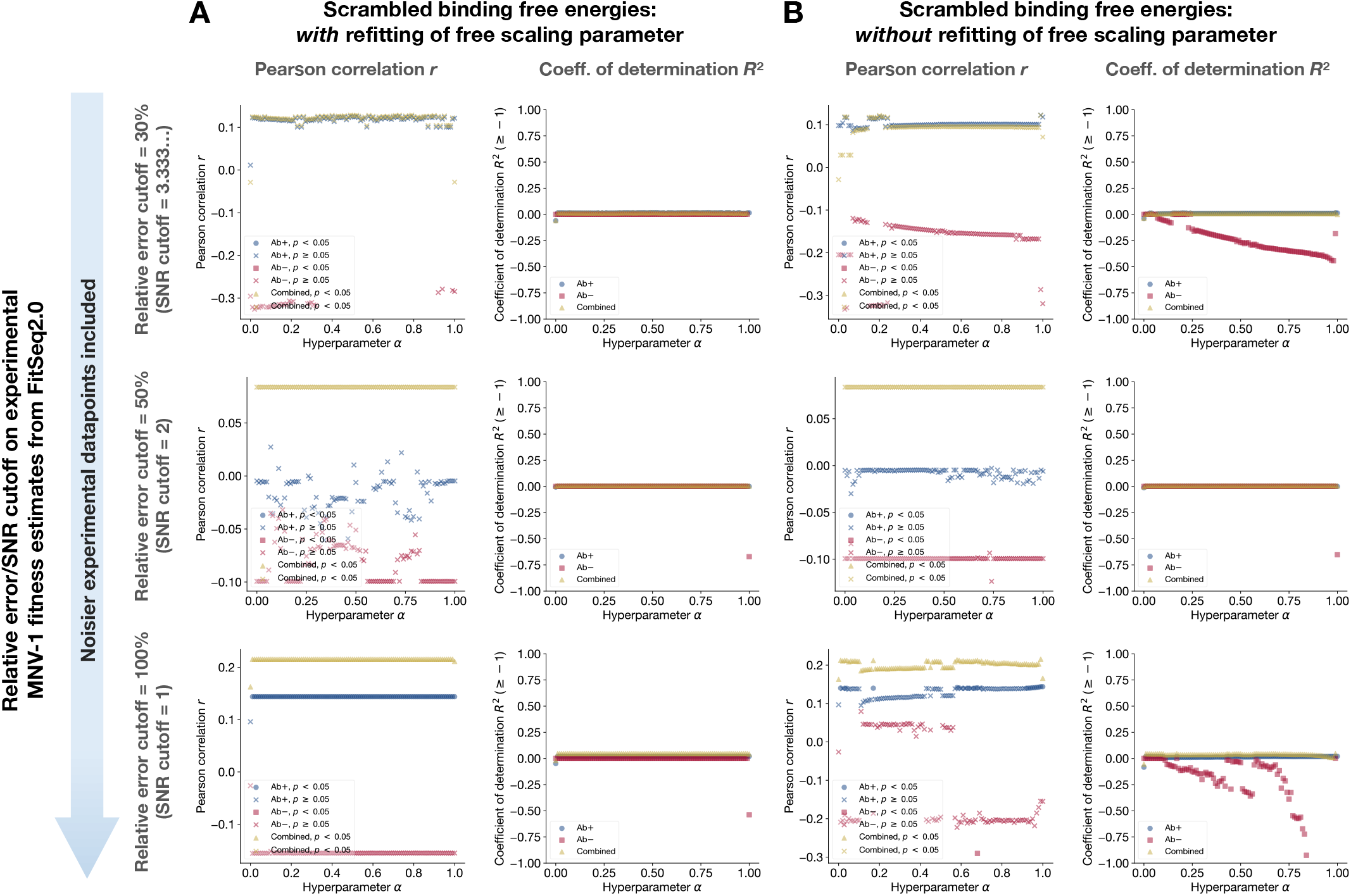
Scrambling EvoEF binding free energy assignments destroys nearly all correlations between fitnesses obtained from Fit-Seq2.0 and fitnesses predicted computationally using the *in vitro* biophysical model with EvoEF binding free energies. For the MNV-1 biophysical model fits, to test robustness of the *in vitro* biophysical model, we independently scrambled the EvoEF-predicted Δ*G*_H_(**s**) and Δ*G*_Ab_(**s**) assignemnts to each strain **s**—for each dataset—and re-performed the fitting to empirical Fit-Seq2.0 data in order to ensure that the predicitve power was not simply due to the number of free parameters in the fitting function. We found that Pearson correlations *r* and coefficients of determination *R*^2^ were nearly destroyed at all relative error cutoffs for both (**A**) the model in which the free multiplicative scaling parameter was refit across Ab+ and Ab− datasets and (**B**) the model in which the free multiplicative scaling parameter was not refit across Ab+ and Ab− datasets. The vast majority of Pearson correlations’ did not have *p*-values below 0.05.

**Figure S4:**
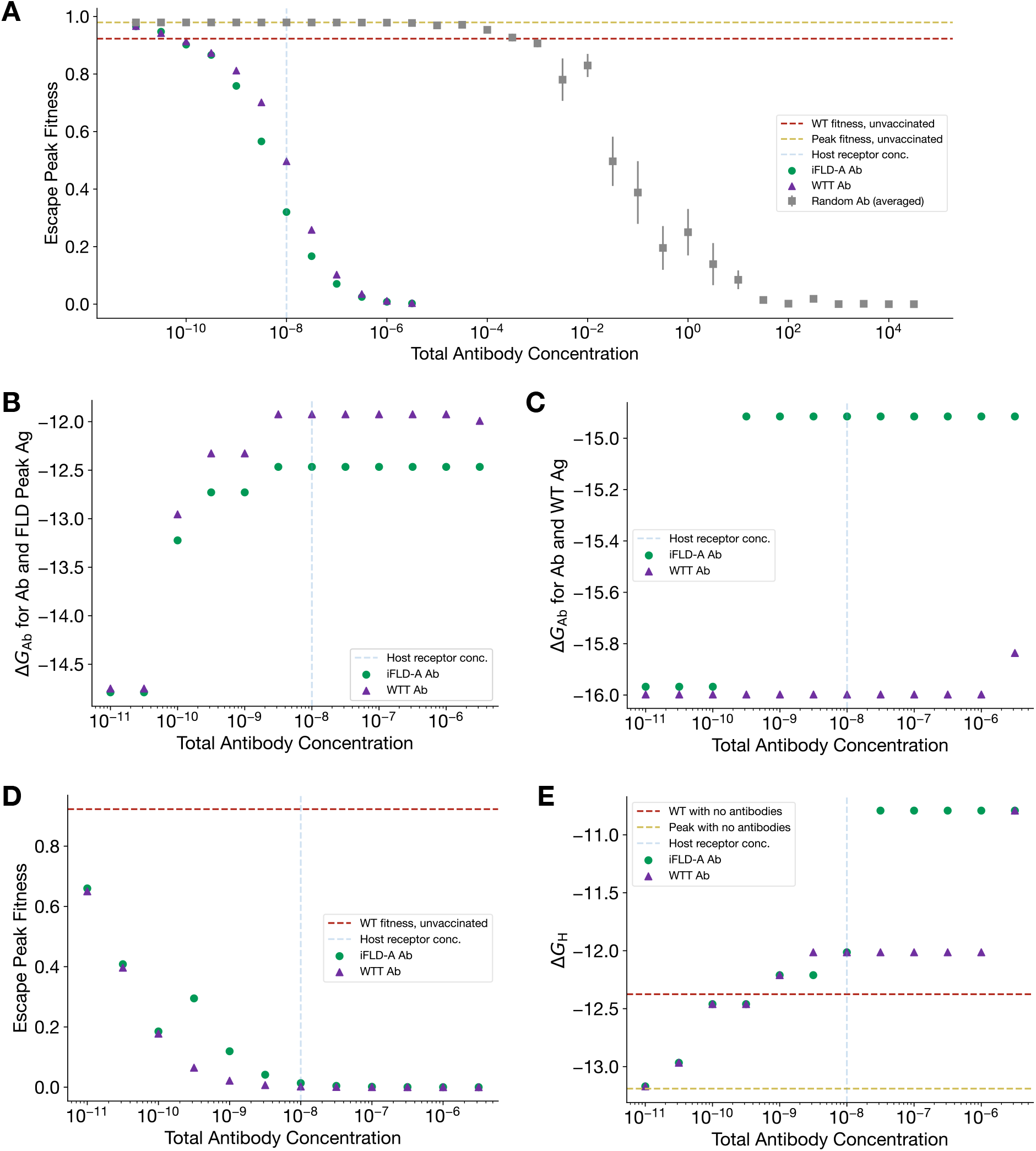
Extended data for iFLD-A protocol with SARS-CoV-2 wildtype. (**A**) Extended range of random antibody concentrations show fitness peak reduction is only possible at concentrations much higher than host concentration. (**B**) The iFLD-A antibody binds tighter to its target antigen than the WTT antibody does. (**C**) The iFLD-A antibody binds less strongly to the wildtype antigen than the WTT antibody does. (**D**) Despite weaker binding to the wildtype antigen, the iFLD-A still substantially suppresses wildtype fitness, often reducing it as much as the WTT antigen. (**E**) Both the iFLD-A antibody and the WTT antibody result in post-vaccination fitness peaks for which the highest-fitness escape variants bind the host receptor less strongly than in absence of any antibody.

**Figure S5:**
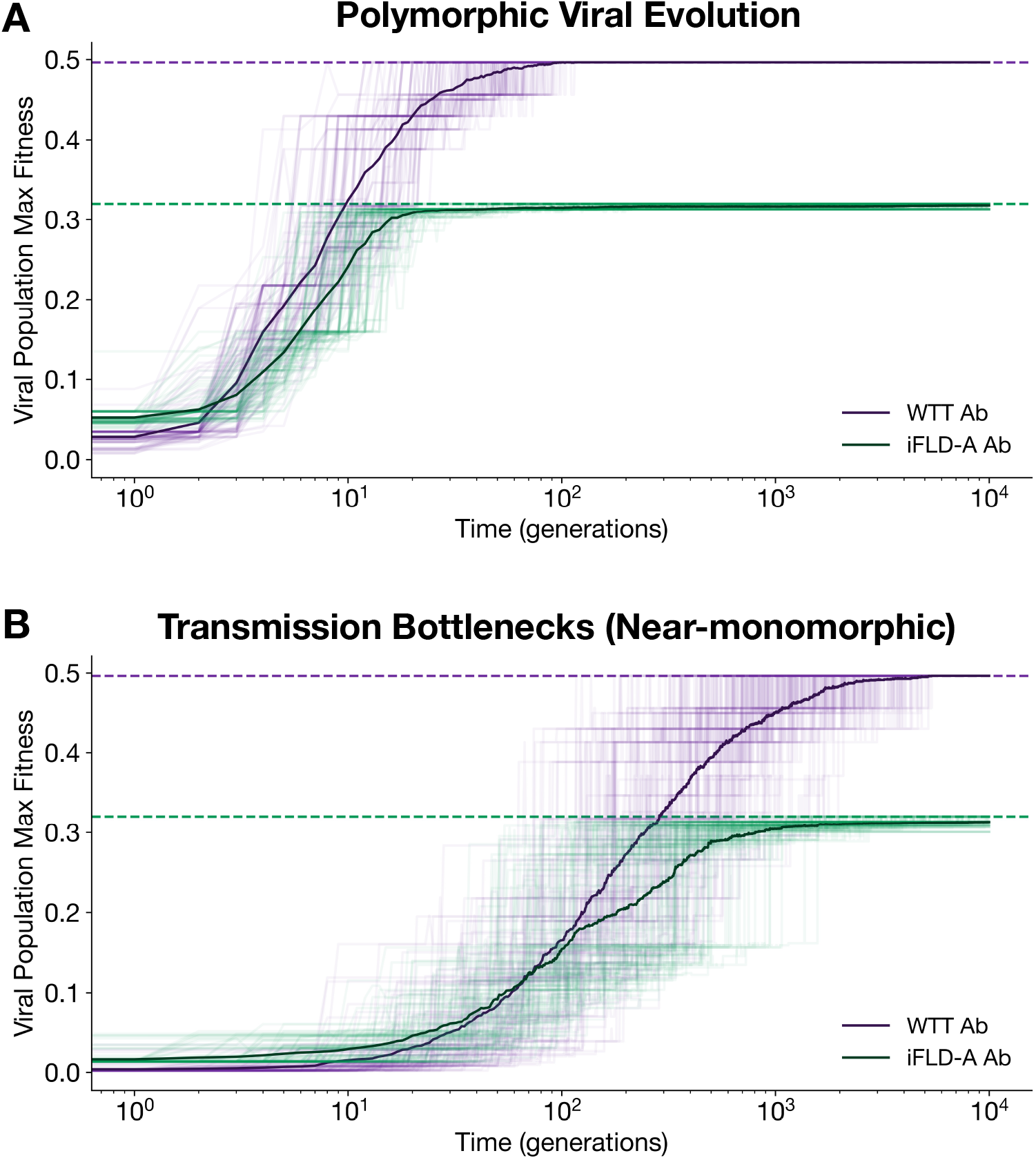
Trajectories of population maximum fitness. (**A**) Polymorphic and (**B**) near-monomorphic viral evolutionary dynamics, with trajectories corresponding to the highest fitness in the population at any given time. In both regimes, populations exhibit slower fitness growth and are trapped under a lower fitness ceiling by the iFLD-A antibody compared to the WTT antibody. Dashed lines indicate global maximum fitness, and translucent lines are population mean fitness trajectories from each of 100 Wright-Fisher trials.

**Figure S6:**
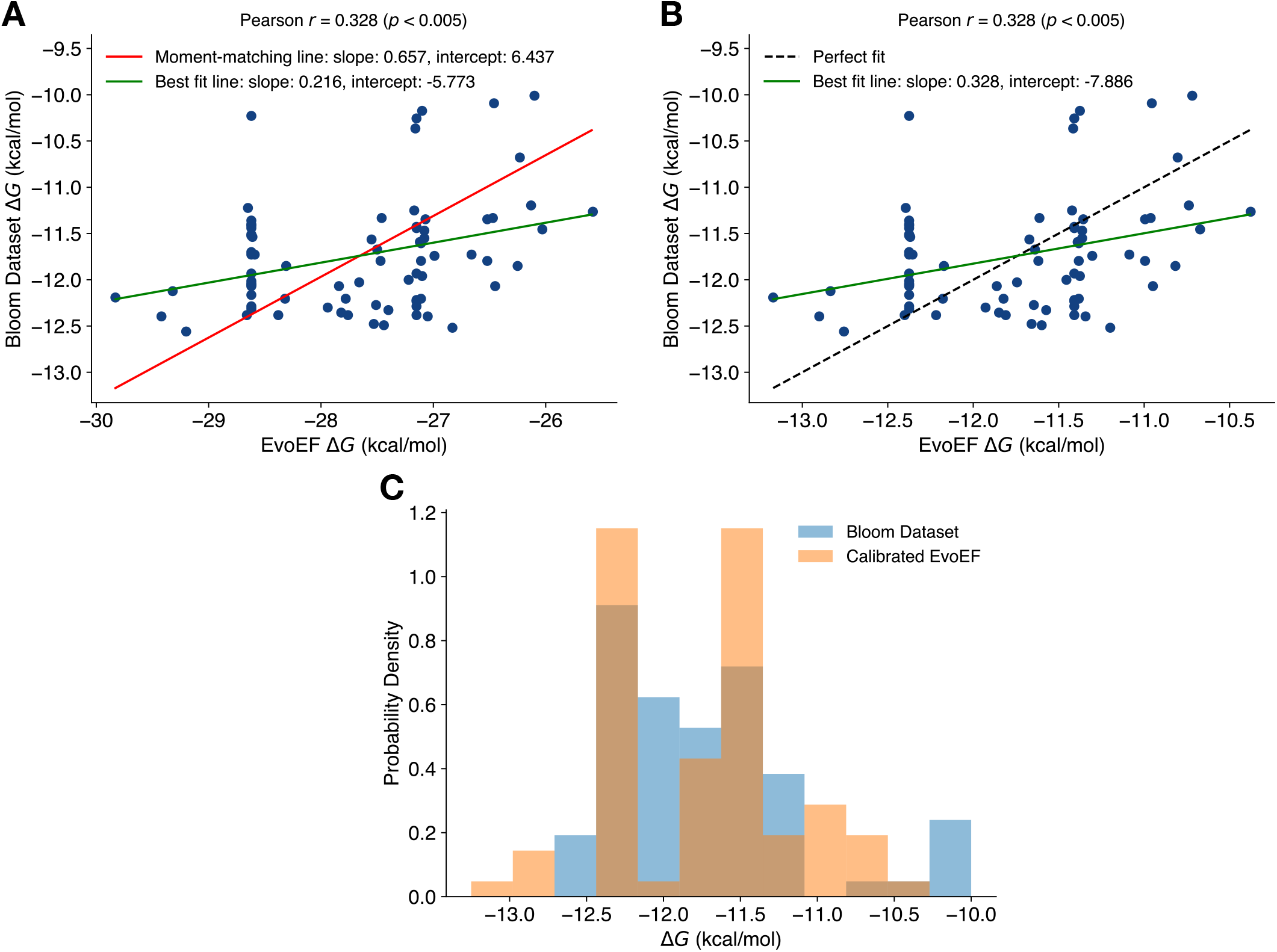
Calibration of host-antigen force field binding affinities to experimental measurements. (**A**) Force field and experimental binding affinities for the 77 antigens present in both datasets are positively correlated. (**B**) Calibration of force field binding affinities using moment-matching helps bring force field binding affinities absolutely closer to experimental ones. (**C**) The experimental data distribution and the calibrated force field data have matching means and variances.

**Figure S7:**
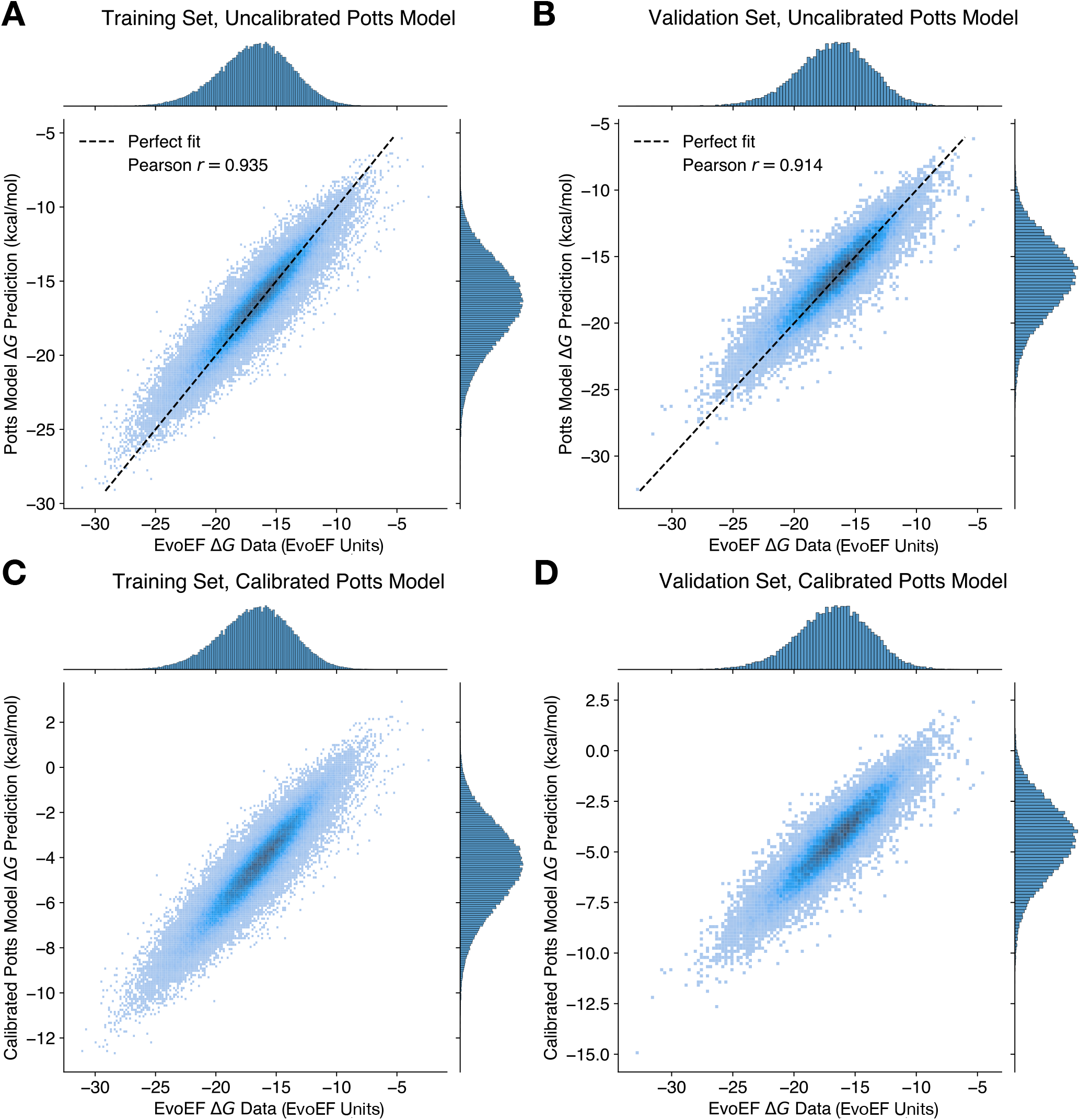
Training, validation, and calibration (to experimental data) of Potts model for antibody-antigen binding affinities. Potts model predictions on (**A**) training and (**B**) validation force field binding affinities show internal consistency. (**C**) and (**D**) show the same, but after calibration of the Potts model to experimental data.

## Notes

### Competing Interest Statement

The authors have declared no competing interest.

### Summary of Updates

In vivo model, experimental and epidemiological data. New figures and text.

